# Physical confinement regulates transition in nematode motility

**DOI:** 10.1101/2025.01.22.633777

**Authors:** Saheli Dey, M Sreepadmanabh, Sayan Kundu, Ashitha B Arun, Sandhya P Koushika, Shashi Thutupalli, Duncan Hewitt, Tapomoy Bhattacharjee

## Abstract

How do worms navigate their complex natural surroundings? Undulatory microswimmers such as nematodes typically inhabit environments such as soil, vegetable matter, and host tissues. While the natural habitats of nematodes are often three-dimensional granular niches with spatiotemporally varying visco-elasto-plastic material properties that impose physical constraints on their motion, current knowledge about nematode motility patterns broadly comes from investigating model organisms such as *Caenorhabditis elegans* either inside liquid cultures or the surface of soft agar pads. How nematodes move through 3D granular niches across different degrees of physical confinement remains poorly understood due to a lack of optically transparent 3D granular matrices. We bridge this gap by engineering an optically transparent granular matrix to directly visualise and quantitatively analyse nematode motion. Importantly, nematodes can freely move through this matrix by generating a minimal yield stress; once the nematode moves away, the matrix self-heals to ensure the material properties remain invariant. Using these platforms, we observe that the propulsive speed of nematodes shows a non-monotonic relation with the yield stress of their microenvironment. This non-monotonicity emerges as nematodes optimize for efficient navigation at higher yield stress, wherein, their forward propulsive speed matches the wave speed along their body. This regulation of locomotory behaviour is purely dictated by the physical interaction of the nematode with its environment without involving soft-touch sensory neurons. Remarkably, predictions from a slender body theory of undulatory motion exactly capture the scaling behaviour for both efficiency and mode of motion as obtained from the experimental data. Finally, in a phase space described by non-dimensional propulsive efficiency and a non-dimensional time scale of motility, we capture a gait transition from poorly efficient thrashing under low confinement to more efficient crawling under high confinement. Thus, our work establishes a new regulatory paradigm describing how distinct modes of undulatory motion emerge under different degrees of physical confinement.

## 1. Introduction

Undulatory motion is a widespread phenomenon in the biological world and is commonly employed by worms, especially nematodes, living in environments such as soil, fruit pulp, and host tissues, to efficiently navigate their complex and disordered surroundings^1,2^. Such niches are also routinely subject to spatiotemporally dynamic agents such as temperature, moisture content, and chemical degradation, which significantly alter their structural properties — forcing inhabitant nematodes to constantly adapt their motility patterns in response to the prevailing mechanical regimes^2^. Understanding such motility patterns has proven instrumental for research on adaptation, ageing, developmental progression, neurological disorders, and mechanosensory responses^3–7^ using model organisms such as *Caenorhabditis elegans*^1^. These nematodes accomplish motion through undulatory contortions of their bodies, which range from thrashing in liquids to surface-level crawling on agar pads^7,8^. However, such observations are largely derived from 2D plates and homogeneously stirred liquids, which only span a subset of the mechanical complexity encountered in natural environments^9,10^. Although past efforts have quantitatively described how altered rheological properties alter undulatory motion, these remain constrained to fluid-like physical regimes^11–18^. Prior attempts at mimicking 2D soft solid-like matrices have employed packings of glass beads, while fruit pulp and paper stacks have been used to generate 3D scaffolds^19–22^. A fundamental limitation of such systems is that even though they allow for burrowing behaviour, the process of nematode motion itself destroys the structural integrity and alters the material properties of these matrices. Furthermore, meaningful biophysical measurements of motility require precise observation and tracking of nematode movement in 3D space—which cannot be afforded by such opaque building materials that preclude direct visualisation (**Fig. 1a**). Consequently, how undulatory microswimmers such as nematodes navigate a truly 3D, granular, and disordered environment across different mechanical regimes has not been systematically explored.

**Figure 1:**
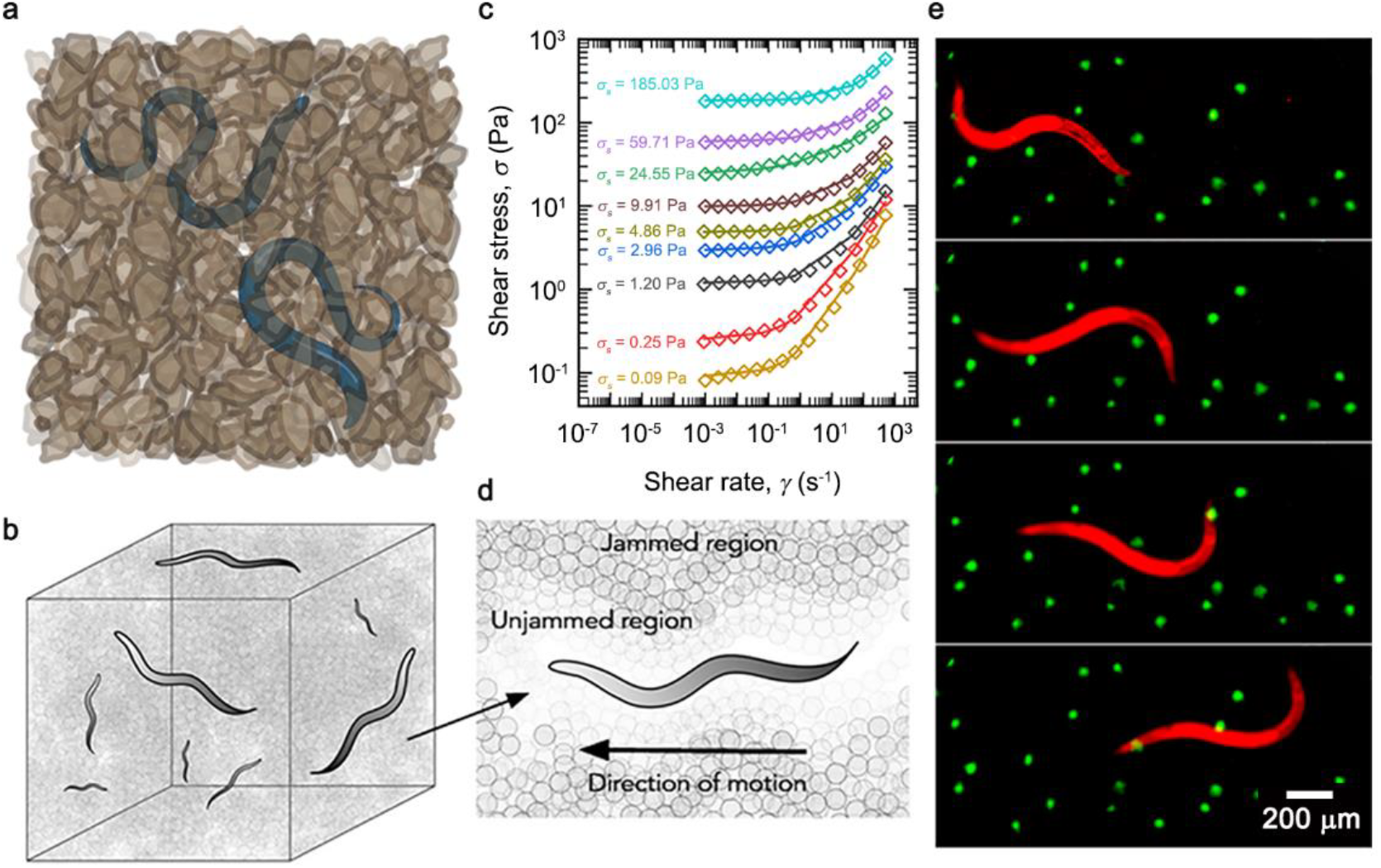
Engineering an optically transparent, mechanically tunable, 3D yield stress matrix for observing undulatory nematodes. (a) The natural habitats of nematodes, such as soil, are granular and porous 3D environments that are also opaque in nature, making it difficult to directly observe and track them in 3D space. (b) Schematic representation of the nematode C. elegans in the optically transparent 3D microgel growth matrix. The matrix comprises of highly-swollen hydrogel granules dispersed in liquid nutrient broth, giving rise to an internally disordered pore-space geometry. Each granule is a randomly crosslinked long-chain polymer with nanometer-scale internal mesh sizes which allow for unimpeded nutrient diffusion throughout the bulk matrix. (c) Characterization of the 3D matrix, as reported using shear rheology. The tunable mechanics of the system enables reproducible engineering of precisely-defined mechanical regimes — representing altered degrees of physical confinement — indicated here by the varying yield stresses of different matrices. (d) The yield stress nature of the system allows nematodes in motion to transiently fluidize the matrix by displacing the hydrogel granules surrounding them, which subsequently recovers in the wake of the nematode motion, thereby conserving the material properties of the system. (e) Nematode motion in a 3D microgel matrix with embedded fluorescent beads, showing that worms only deform their immediate surroundings without causing far-field perturbations — only beads directly in the path of an undulating worm get displaced, whereas beads in the surrounding regions remain at rest.

Interrogating undulatory motion in 3D space requires optically transparent, self-healing, biocompatible, 3D, soft solid-like matrices which stably retain their material properties following repeated rounds of mechanical shear. We address these requirements by introducing an optically transparent, self-healing, viscoelastic granular matrix that presents a porous and disordered 3D microenvironment for nematodes. The material itself comprises jammed packings of hydrogel granules swollen in an M9-based nutrient broth, which can be elastically deformed and reversibly displaced by the nematode’s motion. Importantly, this material exhibits shear-dependent rheology, by transitioning from a soft solid-like state at rest to being locally and reversibly yielded by the nematode’s motion — above a minimal critical stress level termed the yield stress — before immediately recovering the original material properties once the nematode moves away. The highly swollen nature of the individual granules makes the matrix optically transparent due to its low solid fraction, allowing us direct visualisation of nematode motion inside a 3D granular media. Further, we leverage the tunable mechanical properties of this matrix by varying the packing fraction of the hydrogel granules to precisely regulate the degree of physical confinement, which is manifested by an alteration in the material’s yield stress. Tracking nematode motility reveals that the propulsive speed varies non-monotonically with the increase in 3D physical confinement. Combining experimental findings with predictions from slender body theory, we show that the underlying basis for these trends is a maximization of the worm’s forward propulsive efficiency under higher physical confinement.

Across these physical regimes, we observe a progressive transition between gaits, from a thrashing motion in low yield stress regimes — similar to that observed in homogeneous liquid — to a crawling motion in high yield stress regimes — akin to that observed on 2D surfaces. Remarkably, using genetically mutated nematode strains, we find that both the non-monotonic trends in propulsive speed and the optimization of propulsive efficiency are independent of soft-touch mechanosensory responses. Rather, we establish that the rheological properties of the nematode’s microenvironment directly enable context-specific modes of motility. Transforming these results into a phase space defined by the non-dimensionalized efficiency of propulsive motion and a non-dimensionalized time scale between the undulatory frequency of the nematode body and relaxation time of the granular matrix, we observe a transition in nematode gait occurs as higher efficiency of motion emerges with increasing physical confinement. Together, these results explain how nematodes modulate their undulatory patterns to optimize for the most efficient form of forward motion as a function of the material properties of 3D granular media. Our work shows that motion in viscoelastic 3D disordered environments has unexplored facets, capturing and modelling which requires leveraging experimental platforms that allow for direct visualisation of active matter in biomimetic complex mechanical regimes. These findings hold significant implications towards reshaping our ideas of motion in 3D space and suggest important considerations for characterizing the behavioural patterns of undulatory microswimmers. Finally, we view this as an impetus for revisiting our canonical understanding of motility-associated phenotypes as readouts for the regulatory processes controlling development, mechanotransduction, and cognition.

## 2. Results

### 2.1. An optically transparent 3D growth matrix for studying nematode motion

To structurally mimic the complex 3D natural habitats of nematodes, we pack together micron-sized hydrogel particles above jamming concentrations to generate an optically transparent, granular, and porous 3D matrix ^23,24^ (**Fig. 1b**). Each such microparticle comprises of randomly crosslinked long-chain polymers which are highly swollen in the liquid culture media at neutral pH, rendering them optically transparent (**Fig. 1b**). We precisely control the rheological properties of the matrix by altering the solid fraction of the polymer, as evidenced by their tunable shear moduli ^25^. When unperturbed, the material exhibits soft solid-like properties, which are assessed using rheological measurements by applying small (1%) amplitude shear over varying time periods. Herein, we observe that the elastic storage modulus of the material (*G’*) — which denotes an elastic solid-like nature, corresponding to the energy stored within the material while undergoing deformation — is significantly higher than the viscous loss modulus (*G’’*) — which represents the viscous liquid-like nature, corresponding to the energy lost by frictional forces and heat generated during deformation (**SI Fig. 1**). This implies that the material exhibits solid-like behaviour at rest. Importantly, we do not observe any significant change in the *G’* across varying shear frequencies, which demonstrates its time-invariant material properties.

In natural niches such as soil, nematodes locally rearrange the particles during motion, whereas at rest, they remain embedded within a granular microenvironment. This aspect of the nematode habitat is captured by our jammed granular microgel systems; as the microparticles themselves are discrete non-covalently bonded entities, they can be readily displaced by the application of a shearing force which locally yields the material. Importantly, the particles recover this deformation in the wake of the shearing agent, whereby, the mechanical properties of the overall matrix remain conserved despite repeated deforming stresses. To quantitatively characterize this, we subject the matrices to unidirectional shear at varying rates and record the shear stress responses corresponding to each shear rate (**Fig. 1c**). Under high shear rates, we observe a shear-dependent stress response, which subsequently plateaus under low shear rates. This behaviour implies a self-healing property (**SI Fig. 2a**), wherein the 3D granular matrix reversibly transitions from solid-like behaviour under low shear regimes — when the microparticles remain at rest — to fluid-like behaviour under high shear regimes — when the microparticles are transiently displaced (**Fig. 1d**). This crossover from solid-like behaviour to fluid-like behaviour is set by the specific yield stress limit of a given microgel formulation, which determines how much deforming stress the nematode needs to generate to fluidize the surrounding matrix during its movement. To verify this, we observe nematode motion in a 3D microgel bath embedded with homogeneously dispersed fluorescent beads. Indeed, beads embedded in the vicinity of a moving nematode are displaced, whereas beads positioned away from the nematode remain at rest during motion (**Fig. 1e** and **SI Fig. 2b and 2c**). Hence, the undulatory movement only locally and reversibly deforms these matrices, followed by immediate recovery in the wake of a propagating nematode. Furthermore, our microgel growth media does not mechanically deform or compress the nematode bodies, thereby ruling out any experimental artefacts due to aberrant organismal physiology (**SI Fig. 2d**).

**Figure 2:**
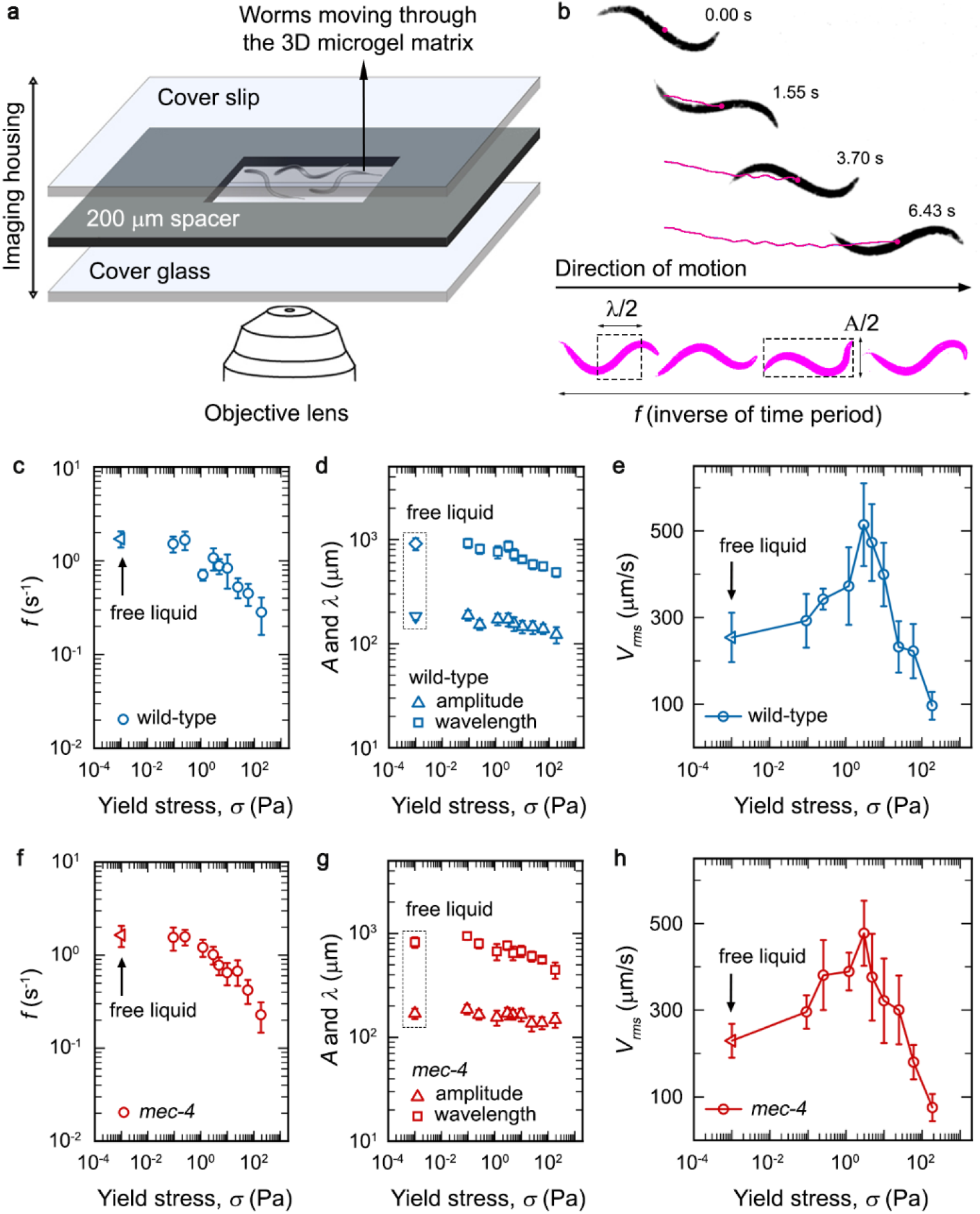
Quantitative analysis of nematode motion reveals a non-monotonic dependence of the forward propulsive speed on matrix yield stress. (a) Schematic representation of the experimental setup for visualizing nematode motion in the 3D microgel matrix. (b) Nematode motion is idealized as a propogating sinusoidal wave, with the specified descriptors of wavelength, frequency, and amplitude of each undulatory motion obtained through quantitative analysis of timelapse imaging performed on individual nematodes. (c and d) While the frequency of undulatory motion shows a progressive decline with increasing matrix yield stresses, both the amplitude and wavelength remain less affected. (e) The forward propulsive speed, *V*_*rms*_, as measured from the centroid displacement of the nematode, shows an unexpected non-monotonic dependency on the matrix yield stress – initially increasing with an increase in physical confinement, followed by a sharp decline under higher degrees of confinement. (f, g, and h) Experiments using soft touch-insensitive mutant *mec-4* strains mirror the trends observed with the wild-type strain, indicating that the observed patterns of yield stress-regulated motility is not specifically driven by this mechanotransduction-based pathway. Since free liquid does not have a yield stress, an arbitrary value of 10^−3^ Pa (viscosity of water times one second) is assigned to include these data on the same plot for panels c-h.

### 2.2. Increasing physical confinement leads to non-monotonic trends in propulsive speed, but not in sideways speed

Leveraging the optically transparent nature of the 3D microgel matrix, we directly visualise and track nematodes embedded within these using a custom-built imaging chamber. We prepare these chambers by placing a 200 μm-deep well — hollowed out from a thick plastic film — sandwiched between a coverslip and a microscopy slide. More precisely, each such imaging well is first placed on a microscopy slide and then filled with the jammed granular microgel matrix, within which a small number of nematodes are carefully embedded. Subsequently, a coverslip is used to seal off this thin slab of the microgel matrix, creating a 3D housing for the nematodes to swim around. Using confocal microscopy, we capture individual nematode motion within this setup as timelapse images at a high frame rate (**Fig. 2a**). Further image processing workflows to threshold and segment the nematode body allow us to quantitatively analyse the nematode motion as a sinusoidal wave propagation (**Fig. 2b**), from which we obtain the components of amplitude (*A*), frequency (*f*), and wavelength of motion (λ). We further use these to determine the sideways speed (*f*· *A*) and wave speed (*f*· λ). Collectively, these parameters allow us to quantitatively describe nematode motion within 3D granular media.

To interrogate whether the undulatory propulsive patterns of nematodes depend on the material properties of their surroundings, we first observe how fundamental descriptors of nematode motion — viz., the frequency, amplitude, and wavelength of individual sinusoidal undulations — are altered across different degrees of physical confinement, represented by varying yield stresses of the 3D matrix. While we observe a progressive decay in the frequency of undulations with increasing yield stress (**Fig. 2c**), both the amplitude and wavelength remain largely conserved across different mechanical regimes (**Fig. 2d**). This suggests that elevated physical confinement has a more pronounced effect on the rate at which the nematode executes undulations, as opposed to the nature of these undulatory patterns. Hence, it is likely that a nematode’s propulsive speed through 3D granular media is strongly influenced by environmental mechanics. To test this, we first quantify the nematode’s overall displacement by tracking the body centroid to obtain its mean square displacement (**SI Fig. 3a**). From the mean squared displacements, we measure root mean square speed (*V*_*rms*_) as a function of lag time (*τ*) which represents the forward propulsive speed (**SI Fig. 3b**). Next, to fix an appropriate time scale for comparing *V*_*rms*_ in different matrices, we analyze the nematode body undulations to identify characteristic frequencies and time periods of motion. We compare the *V*_*rms*_ at lag times corresponding to the time period of motion in a given matrix. Extending this analysis over different regimes of physical confinement — defined by the yield stress of the system — we find that the *V*_*rms*_ shows a non-monotonic dependence on the yield stress of the system (**Fig. 2e**). Surprisingly, an increase in confinement appears to provide a propulsive advantage to these undulatory swimmers up till a threshold yield stress, beyond which, the expected effects of confinement in reducing the speed of motion manifest.

**Figure 3:**
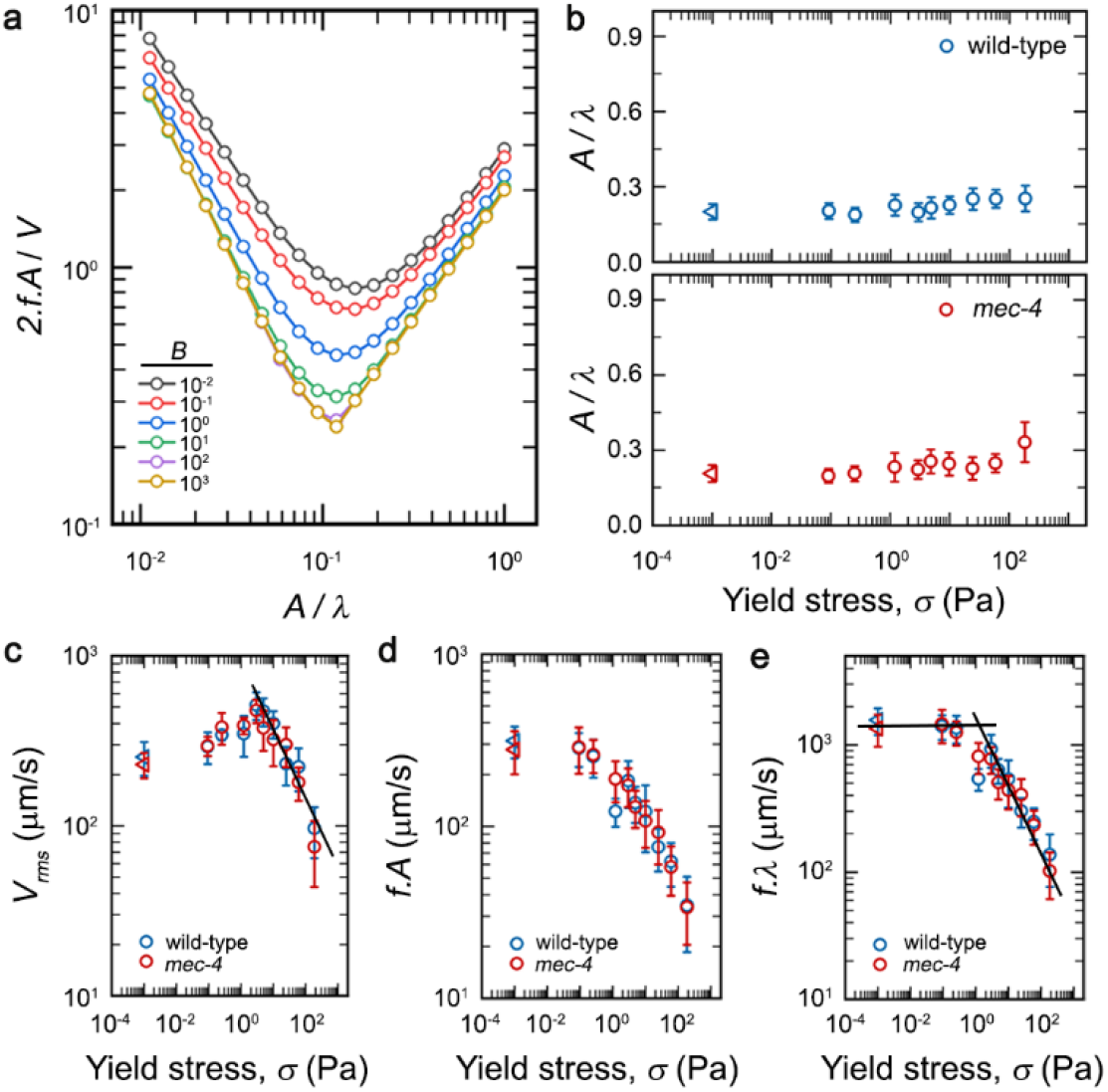
The undulatory motion of nematodes in high yield stress regimes is characterized by small *A*/*λ* ratios as well as a power law dependency of the propulsive and wave speed on yield stress. (a) Numerical solutions comparing the Strouhal number against different body forms (*A*/*λ* ratio) show a generalized non-monotonic behaviour across different mechanical regimes (varying *B*). (b) Experiments reveal that nematodes maintain their body form (*A*/*λ* ratio) at a largely constant and small magnitude (*A*/*λ* ≤ 1), as predicted by the theoretical model, across different degrees of 3D physical confinement. (c) Pooled datasets of the propulsive speed (*V*_*rms*_) show a distinct power law dependence on the matrix yield stress under higher degrees of physical confinement, which decays as 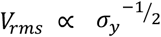. (d) The sideways component of motion (*f* · A) undergoes a progressive and sustained decrease with increasing physical confinement. (e) The wave speed (*f* · λ) of the undulatory motion remains initially constant, before exhibiting a decay with increasing yield stress, which follows a scaling behavior similar to that of the propulsive speed, given by 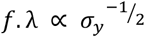. The experimentally-observed scaling behavior of both *V*_*rms*_ and *f* · λ are in good agreement with predictions from the theoretical model. In all the plots of panel b, c, d,and e, the triangular symbols represent data from nematodes moving through free liquid. Since free liquid does not have a yield stress, an arbitrary value of 10^−3^ Pa (viscosity of water times one second) is assigned to include these data on the same plot.

In our experiments, the nematode moves through a bath of granular particles. Hence, it is conceivable that its motion is regulated by physical interaction with the particles and touch sensation, which actively effect behavioural changes leading up to an alteration in the nematode motion. We test this hypothesis by employing a gentle-touch sensory mutant (*mec-4*), which carries a genetic aberration rendering it insensitive to soft mechanical pressures^26^. Given the low shear moduli of our microgels in comparison to 2D agar/plastic/glass substrates, gentle-touch mechanosensation represents a putative module for environmental perception by nematodes moving in granular 3D media. Surprisingly, we find no significant differences between the wild-type and *mec-4* mutant strains for any of the descriptors of undulatory motion, across the entire range of physical confinement interrogated by our study (**Fig. 2f** and **2g**). Indeed, the *V*_*rms*_ of soft touch-insensitive mutant nematodes also exhibit a non-monotonic dependence on yield stress, mirroring the trend observed for the wild-type strain (**Fig. 2h**). Furthermore, the non-monotonic nature of the forward propulsive speed across different mechanical regimes remains robustly conserved for both wild-type and *mec-4* strains, irrespective of the time period of motion (**SI Fig. 3c and 3d**). Together, these observations suggest that the mechanism via which the forward propulsive speed of nematodes is modulated in a yield stress-dependent manner is likely not driven by active neuronal perception and consequent biochemical response to altered mechanical regimes. Rather, we hypothesize that such trends reflect how 3D confinement effects a physical regulation of undulatory motion.

### 2.3. A slender body theory-inspired formulation predicts the dependence of wave speed on yield stress

To quantitatively understand how different degrees of physical confinement imposed by the 3D microgel matrix regulate nematode motion, we use predictions from a slender body theory to describe the interactions involved in this process. We adopt a previously described formulation^27^, to conceptualize nematode motion through a yield stress matrix, which can be modelled as a Herschel-Bulkley fluid obeying the following relation

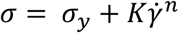

for shear stress σ, yield stress σ_*y*_, consistency *K*, index *n*, and the strain rate 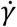. We consider *n* = 0.5 throughout, which is in good agreement with our experimental medium. Further, we incorporate the sinusoidal nature of the worm body undulations by modelling its propagation using the idealised waveform

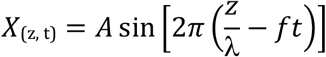

Given such a waveform, a resistive-force theory approach allows us to determine the forward speed *V* and power input *P* as functions of the waveform (i.e. *A, λ* and *f*), and the rheology of the matrix. Our approach^27,28^ relies on the assumption that deformation is relatively localised to the long and thin organism (details are provided in **SI section 2**). The importance of the yield stress can be characterised by the dimensionless Bingham number, *B*, as

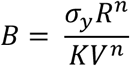

Given a value of *B* and a body form — defined as the ratio between amplitude and wavelength (*A*/*λ*) — our theoretical formulation returns the forward speed *V* and the power *P*. A simple measure of efficiency is provided by the Strouhal number — defined as the ratio between the sideways component of motion (*f*· *A*) and forward speed (*V*) — the magnitude of which inversely correlates with the efficiency of forward motion with respect to the lateral displacement. Interestingly, we find a non-monotonic behaviour in the Strouhal number as a function of the body form (*A*/*λ*), which, along with a screening for the optimal *A*/*λ* ratio across different regimes of *B*, implies that optimal conditions favouring forward motion occur within a range of 0.1 ≤ *A*/*λ* < 0.5 (**Fig. 3a and SI Fig. 4a**). Together, these results indicate that efficient propulsive behaviour in yield stress environments representative of our experimental system requires a pattern of undulatory motion consisting of comparatively small amplitudes — indicative of sideways displacement — relative to the wavelength of undulation — indicative of forward displacement — to achieve efficient forward motion while minimizing lateral movement. Furthermore, we test whether the key assumptions underlying this formulation remain conserved within our experimental paradigm. Encouragingly, for both wild-type and soft-touch insensitive mutant *mec-4* strains, we observe that in high yield stress regimes, the body form (*A*/*λ*) for the nematode’s undulatory motion remains both largely constant and small, falling within the 0.1 ≤ *A*/*λ* < 0.5 range, which mirrors the exact constraints necessitated by the theoretical model (**Fig. 3b**).

**Figure 4:**
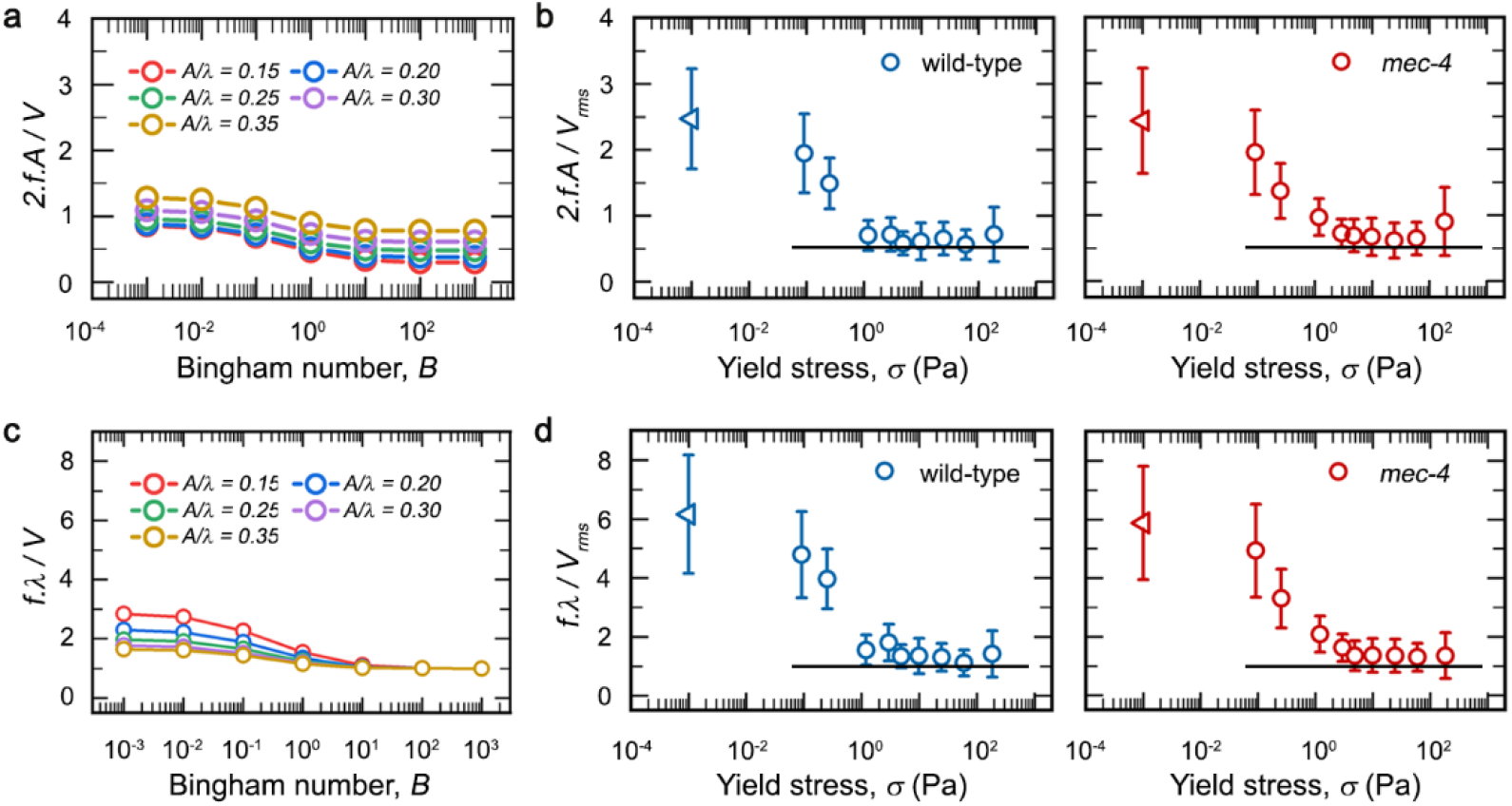
Nematodes optimize their motion in response to elevated physical confinement by minimizing sideways motion, whereby the wave speed along their body matches their forward propulsive speed. (a) Numerical analysis shows that the Strouhal number attains a plateauing behaviour under solid-like regimes (*B* > 1) for all different body forms(*A*/*λ*). (b) Under high yield stress regimes, both wild-type and *mec-4* soft-touch mechanosensory mutant strains optimize for effective forward propulsion by minimizing their sideways component of motion (*f.A*) in comparison to their forward propulsive speed (*V*_*rms*_). (c) Under solid-like mechanical regimes, numerical analysis shows that nematodes match their wave speed (*f*·*λ*) to their forward speed (*V*), across all different body forms. (d) Within high yield stress matrices, the wave speed along nematode bodies matches their propulsive speed, for both wild-type and *mec-4* strains. In all the plots of panel b and d, the triangular symbols represent data from nematodes moving through free liquid. Since free liquid does not have a yield stress, an arbitrary value of 10^−3^ Pa (viscosity of water times one second) is assigned to include these data on the same plot.

Next, we plot all experimental propulsive speed (*V*_*rms*_) data from both the wild-type and soft touch-insensitive mutant strains and observe that the speeds of nematodes follow a universal scaling with the yield stress of the matrix. Precisely, in high yield stress regimes, the propulsive speed shows a power law dependence on the yield stress (**Fig. 3c**). We find that with increasing yield stress, the *V*_*rms*_ decays with a power law exponent of the order of approximately −1/2, such that

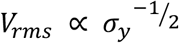

Interestingly, however, the non-monotonic trend holds true only for the forward propulsive speed, *V*_*rms*_. A comparison of the sideways component of motion, given by the product of frequency and amplitude (*f*· *A*), instead shows a progressive decline with an increase in the yield stress of the 3D matrix, even under regimes of low physical confinement (**Fig. 3d**). This contrast between both the *V*_*rms*_ and the sideways speed indicates that even within a small range of increase in yield stress, motion within the 3D matrix manifests distinctly different effects on forward propulsion and sideways motion; while the former shows a transient increase followed by a sustained decline, the latter almost immediately undergoes a gradual decrease with increasing yield stress.

The theoretical formulation predicts a relationship between the waveform parameters *A, λ*, and *f*, the forward speed (*V*), and the power (*P*) expended by the nematode. In general, there is no analytical expression for this relationship, but for amplitudes in the optimum range outlined above and for large enough yield stress, previous reports have arrived at numerical solutions as ^27,28^ (elaborated in **SI Section 2**)

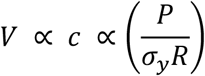

where *c*, given by *c* = *f*· *λ* represents the wave speed along the worm body.

Thus, the experimental observation that the propulsive speed (*V*_*rms*_) decays as the square root of the yield stress immediately implies that the power is increasing as 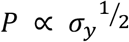, and the wave speed of motion in the solid-like mechanical regime (*B* >1) is

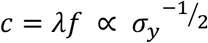

Remarkably, we find that our experimentally measured data on the wave speed (*f*· l) of nematodes across different mechanical regimes also exhibits a similar scaling behaviour under high yield stress (**Fig. 3e**). This strong agreement between the theoretical prediction and the experimental data clearly indicates that the observed modulation of nematode motility across different degrees of physical confinement is dependent on the matrix yield stress properties. Furthermore, the correspondence between both wild-type and soft touch-insensitive mutant nematodes underscores that the phenomena reported herein are driven by generalizable interactions between the worm motion and their environment, rather than touch-specific neuronal signalling.

Interestingly, it might seem plausible that the organism exerts a fixed amount of power *P*, which, given the scaling reported above, would seem to suggest that the locomotion speed and wave speed should decay like 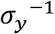 for large yield stress; a somewhat stronger scaling than we observe here. However, this theoretical scaling for the power is in practice only attained for very large *B*^28^, and it may simply be that the experiments here are yet to attain this high yield stress limit.

### 2.4. Nematodes optimize their forward propulsive efficiency under higher confinement, wherein wave speed matches the propulsive speed

Following the observations from our theoretical framework and experimental findings above, we next ask how nematode motion is mechanistically regulated to accomplish efficient forward propulsion under high yield stress regimes. To elucidate this, we first numerically solve for the efficiency of forward motion, as captured by the Strouhal number, at different mechanical regimes represented by varying *B*. We find that even small variations in the body form (*A*/*λ* ratio) influence efficiency of forward motion (**Fig. 4a**). As expected, the overall efficiency comes out to be inversely dependent on the *A*/*λ* ratio, since the Strouhal number is directly proportional to amplitude. Interestingly, however, beyond *B* > 1, which indicates a solid-like regime (since the yield stress component outweighs the viscous shear applied by the nematode), the efficiency of forward motion plateaus regardless of the body form, suggesting that a more solid-like mechanical regime enables the conservation of the forward propulsive efficiency. We further validate this by considering the Lighthill efficiency (**SI Fig. 4b**), which gives the relation between the movement speed and expended power — the higher the index, the more effective the nematode’s motion. Strikingly, our numerical results predict that in solid-like mechanical regimes (*B* > 1), Lighthill efficiency also becomes independent of Bingham number, indicating that peak propulsive efficiency is both enabled and constrained by elevated physical confinement (**SI Fig. 4b**). Together, our findings indicate that a decrease in the Strouhal number captures an increase in propulsive efficiency as depicted by the Lighthill parameter. Hence, from here on, we will consider the Strouhal number as the metric for the efficiency of propulsive motion in complex environments.

Interestingly, the experimental measurement of the Strouhal number for wild-type as well as *mec-4* mutants across different degrees of confinement both quantitatively and qualitatively captures the theoretical predictions. More precisely, the experimental Strouhal number shows an initial decline with an increase in yield stress, followed by a subsequent stable plateau, akin to the numerical results (**Fig. 4b**). We also note that the average body form ⟨*A*/*λ*⟩ in our experiments remains largely constant, with a value of approximately 0.25. Remarkably, for this regime, our theoretical predictions of Strouhal number match identically with our experimental findings for both the wild-type and *mec-4* strains (indicated by the guidelines in **Fig 4b**) which shows that propulsive efficiency is directly regulated by the interaction between microenvironment mechanics and bodily undulations. Interestingly, the transition point where the stable Strouhal number regime commences matches the yield stress that gives the maximal *V*_*rms*_. This experimental regime corresponds to a Bingham number of *O*(1) (**SI Fig. 5**), which in turn mirrors trends in the numerical analysis observed when *B* > 1. Together, these results solidify our hypothesis that nematodes optimize their motion to achieve maximal forward propagation while undergoing minimal sideways displacement upon an increase in the degree of physical confinement.

**Figure 5:**
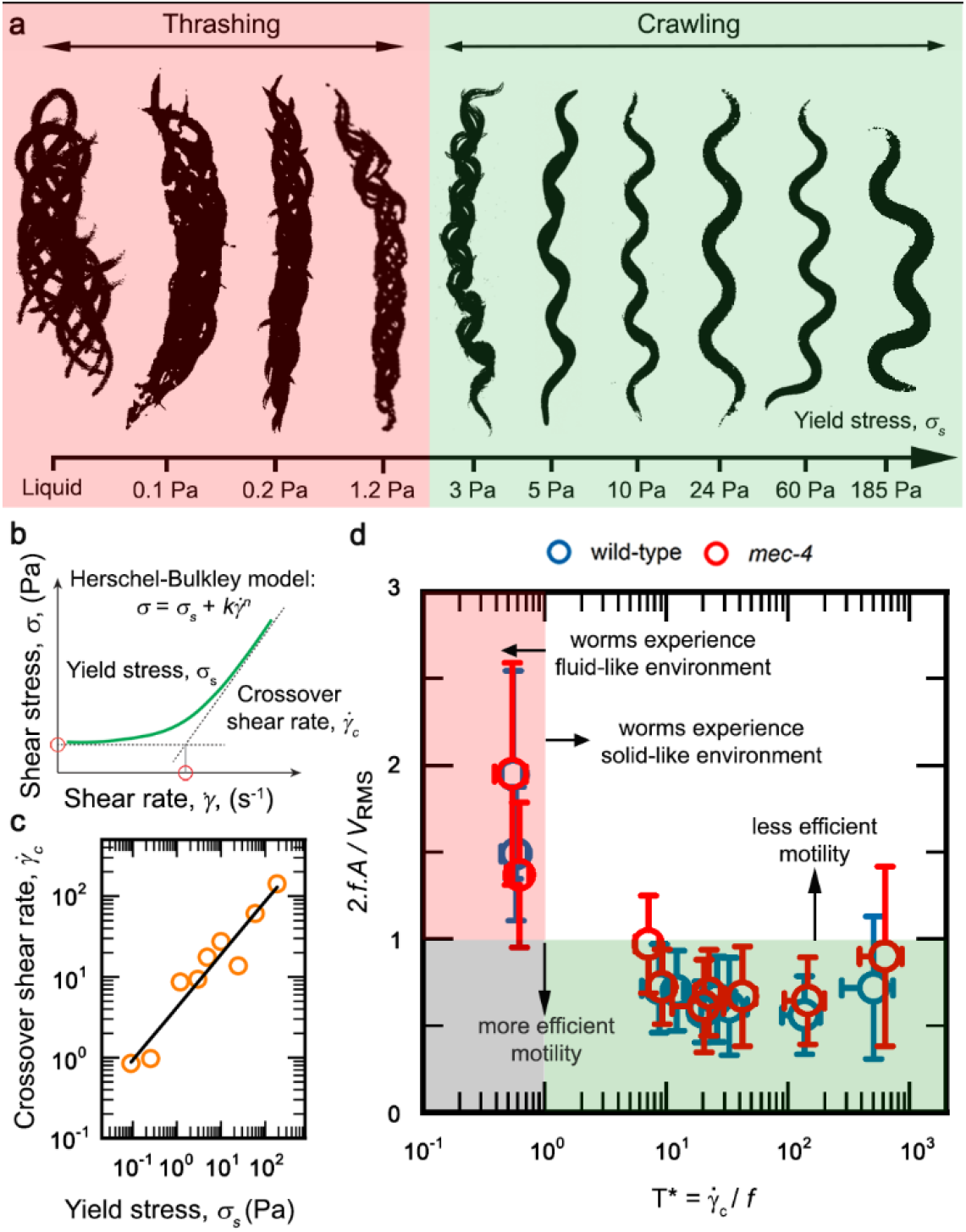
Nematodes exhibit a progressive gait transition in response to increasing physical confinement, which underlies the optimization for forward motion in high yield stress regimes. (a) The undulatory motion of nematodes in the 3D matrix shows a progressive gait transition from thrashing-like behaviour in fluid-like regimes to crawling-like motion in more solid-like regimes. (b) The yield stress nature of the 3D microgel matrix follows a Herschel-Bulkley fluid-like behaviour, showing shear-dependent rheological properties. (c) Each such matrix is characterized by a specific crossover shear rate 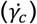, which indicates the time scale of elastic recovery following shear-induced solid to liquid transition — higher 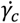 indicates faster recovery of solid-like elastic properties, and vice versa. (d) A 2D phase space defined between the Strouhal number (indicating the efficiency of forward motion) and *T*^∗^ (a measure of the matrix recovery time under nematode motion-induced shear, relative to the rate of nematode undulations) shows two distinct regions: in more fluid-like environments (low *T*^∗^), nematodes exhibit inefficient forward motion (high Strouhal number value), whereas in more solid-like environments (high *T*^∗^), nematodes increase the efficiency of forward motion (low Strouhal number value).

Next, we employ the theoretical model of worm motion to understand how the ratio between the wave speed (*f*· *λ*) and the forward speed (*V*) depends on the environmental mechanics (*B*). We find that as the mechanical regime becomes increasingly solid-like (*B* > 1), the forward speed of the nematode does appear to match its wave speed, regardless of the body form (*A*/*λ*) — exactly as predicted by our model above (**Fig. 4c**). We next examine the experimental data and find that the ratio between the wave speed and the propulsive speed also shows an initial decline, followed by plateauing at critical yield stress, beyond which both remain approximately equal — perfectly mirroring our theoretical predictions (**Fig. 4d**). Similar to the motility efficiency as captured by the Strouhal number, onset of this transition occurs at the exact yield stress wherein the propulsive speed (*V*_*rms*_) is maximal and the Bingham number is of the *O*(1) (**SI Fig. 5**). For additional consistency, we also plot the experimentally derived parameters describing nematode motion against the experimentally determined Bingham number and observe similar trends (**SI Fig. 5** and **6**). Hence, we infer that nematodes optimise their forward propulsive efficiency under elevated physical confinement when the forward propulsive speed matches the wave speed along their body.

### 2.5. Increasing physical confinement alters nematode gait from thrashing to crawling, making forward propulsive motion more efficient in solid-like regimes

Our present consensus on *C. elegans* motility demarcates two distinct forms of motion, varying from a thrashing motion in bulk fluids to crawling on 2D surfaces. However, these models do not clearly account for how physical confinement within granular 3D materials alters the propulsive behaviour. Further, while past work using different species of worms has looked at burrowing behaviour within sediment beds^29,30^, our understanding of how different degrees of physical confinement enable or hamper specific gaits of motion (viz. thrashing or crawling) remains incomplete. Here, considering the stark differences between the trends in propulsive speed and sideways speed with increasing environmental yield stress, we hypothesize that confinement-induced alteration to the nematode’s motion might help explain these patterns. To test this, we generate time projections of nematode motion for identical durations across different degrees of physical confinement (**Fig. 5a**). Comparison between these reveals a stark difference testifying to our quantitative analysis: within low yield stress materials, nematodes execute a thrashing motion, while gradually transitioning to a crawling motion with an increase in yield stress, which is set by the degree of 3D confinement. These observations also fall in line with our measurements of nematode body motion; while the sideways speed — maximal when purely thrashing, minimal when purely crawling — decreases with increasing yield stress, the propulsive speed becomes adjusted to the wave speed along the nematode body (as enabled by crawling behaviour) under higher degrees of confinement. Together, these observations underscore that the physical confinement-driven regulation of nematode motility is dependent on the gait transitions induced by the yield stress of the 3D matrix.

This begs the question — why exactly is the yield stress property of the material influencing nematode gait and forward propagation? Using shear rheology, we previously observed that our 3D growth matrices show yield stress behaviour, transitioning from a viscous fluid-like behaviour under high shear rate to a more elastic solid-like behaviour under low shear. The transition between these regimes is characterized by the yield stress and its corresponding crossover shear rate, which shows a correlated behaviour (**Fig. 5b** and **5c**). Specifically, the crossover shear rate represents the shear rate at which both the viscous and elastic component of the the shear stress are equal, representing the solid to liquid transition. This implies that higher confinement matrices recover faster from mechanical shear as opposed to low confinement matrices. Since nematodes moving through the 3D matrix also generate deforming shear stresses on the material, we directly compare the timescales of the nematode-generated stress against the matrix crossover shear rates to better understand the microenvironmental mechanics in the vicinity of a moving nematode. For this, we define *T*^∗^, as the ratio between the matrix crossover shear rate and the frequency of nematode undulation, as

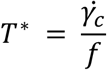

Effectively, *T*^∗^ < 1 implies that the nematode body undulates faster than the matrix relaxation time, which makes the nematode experience a fluid-like environment. By contrast, *T*^∗^ > 1 makes the nematode experience a solid-like environment since the nematode undulation is slower than the material relaxation time. To quantitatively assess if the variation in matrix properties leads to a transition in motion, we define a phase space described by non-dimensional efficiency (Strouhal number) and a non-dimensional time scale of motility (*T*^∗^). Herein, we capture four distinct zones: (I) the red zone, wherein highly undulatory motions (akin to thrashing) give rise to less efficient motility under low confinement; (II) the grey zone, wherein we would hypothetically expect efficient propulsion under low confinement; (III) the white zone, wherein we would hypothetically expect poorly efficient propulsion under high confinement; and finally, (IV) the green zone, wherein crawling-like behaviour enables highly efficient motion under high confinement (**Fig. 5d**). Transforming our experimental findings into this phase space reveals that our data lies squarely within the (I) red zone and the (IV) green zone. Importantly, we observe a sharp transition between the zones when the nematode’s propulsive efficiency increases, which also coincides with the gait transition from thrashing to crawling. It is noteworthy that this occurs within a narrow range of increasing matrix yield stresses — going from just about ~1.2 Pa to ~3 Pa during this transition. Collectively, these analyses quantitatively capture the transition from a more undulatory and less efficient motion (red zone) to a less undulatory and more efficient motion (green zone) with an increase in the degree of physical confinement. This is also validated by the fact that a comparison between *T*^∗^ and sideways speed shows a monotonic decay in the sideways locomotory component with increasing *T*^∗^ (**SI Fig. 7**). Together, these phase spaces describing nematode motility beautifully capture a crossover from thrashing to crawling behaviour as a function of the physical confinement, whereby worms optimize their motion for greater forward propulsive efficiency within increasingly solid-like environments.

## 3. Discussion

Our work introduces the first-ever experimental platform to directly observe, track, and measure nematode motility within an optically transparent, mechanically tunable, viscoelastic, 3D granular matrix. We report that under increasing degrees of physical confinement, forward propulsion shows a non-monotonic trend, whereas, sideways motion monotonically decreases. Importantly, the nematodes undertake these alterations to optimize their efficiency of forward propulsion under increasing confinement, which is achieved when their propulsive speed matches the wave speed along their body. These effects manifest in the form of a gait change from thrashing behaviour in low yield stress regimes to crawling within higher yield stress matrices. Interestingly, in a phase space between propulsive efficiency and a dimensionless time scale of motion, *T*^∗^ — set by the ratio between matrix relaxation time and nematode body undulation frequency — the motility transitions between two distinct regimes: one, wherein inefficient forward motion dominates under fluid-like environments, and two, wherein increasingly efficient forward motion correlates with increasingly solid-like environments. Our work hence establishes a mechanistic basis for the regulation of undulatory motility by 3D physical confinement.

Specifically, in their natural settings, undulatory microswimmers traverse a diverse range of mechanically complex habitats which are granular 3D environments. It is conceivable that sensory perception and subsequent responses within these milieus are governed by a specialized set of neuronal networks and associated biochemical signalling modules. To date, these have remained largely unidentified due to technical limitations precluding direct observation of worm motion within mechanically tunable 3D systems. The unique structural properties of the 3D granular microgel-based platform introduced in this work position it as an excellent candidate for the identification of such putative biological effectors. Parallely, behavioural responses to chemotactic cues are widely used as a readout of neuronal activity and physiological functionality. However, past work on this typically employs flat agar pads with constrained two degrees of freedom. It is an intriguing prospect to explore how both chemoattractant/repellent gradients in 3D alter detection and consequent responses, and how the behavioural changes induced by chemical signalling co-ordinately regulate the nematode’s motion alongside different degrees of physical confinement — effectively, a mechano-chemical regulatory framework of worm motion. This is particularly intriguing, given our observation that nematode gait is directly influenced by the matrix yield stress. This also has broader implications for studying activities such as foraging for food (based on chemotactic detection of bacterial prey) and collectively coordinated motion by strains exhibiting social behaviour. Moreover, this sets up the stage for exploring how genetically encoded behavioural patterns in nematodes can undergo orthogonal modulation by the mechanical properties of their microenvironment.

In general, such a conceptualization is not without precedent. Indeed, past work with short-term bodily conformations of crawling nematodes has used a dynamical systems approach to analyze the resultant eigenworm data across time, yielding predictive phase spaces of worm behavior^31^. A crucial insight here is that the state of a nematode’s body determines not only its locomotory patterns but also the observed variability therein. This implication — that body mechanics capture behavioural patterns^32^ — strongly echoes through our present work as well, albeit with a key difference. While past work has been restricted to data from 2D surfaces or homogeneous liquid media, our exploration of a granular, viscoelastic 3D environment provides new insights into how yield stress properties and material elasticity govern bodily gait and locomotory efficiency. Bridging these seemingly separate frameworks builds up to a fundamental question — are previously reported phase spaces of nematode behaviour distinct from their state in 3D space, or, do both reside within a universal paradigm of behavioural regulation mediated by bodily conformations and microenvironmental mechanics? Resolving this has profound implications for how we seek to model dynamic biological activity in complex environments.

Our present work hence extends two key advancements, both technical and conceptual. The experimental platform introduced here will be of broad interest to the worm community, by enabling the direct interrogation of nematodes in tunable biomimetic 3D settings. Further, we have identified a mechanistic basis for the biophysical regulation of motility, purely based on environmental constraints. This opens up future opportunities for exploring the axes between genetic regulation, behavioural responses, and microenvironment mechanics-induced feedback.

## 4. Materials and Methods

### 4.1. Preparation of 3D growth matrix

We manufacture optically transparent 3D growth matrices by dispersing granules of Carbomer-980 (Ashland Inc.) in liquid M9 nutrient broth. Following homogeneous dissolution of the granules, we adjust the pH of the suspension to neutral pH 7 using 10N NaOH, which causes the hydrogels to significantly swell and form a jammed 3D packing ^33^. The low solid fraction used to manufacture these microgels (between 0.6%-5% of the total weight) ensures that the system is highly optically transparent. By altering the packing fraction and degree of swelling of the microgel granules, we exercise precise control over the rheological properties of the 3D matrix ^34^. We further characterize the viscoelastic properties of these matrices as described below to represent different degrees of confinement.

### 4.2. Rheological characterization

Using shear rheology performed on an Anton Paar MCR302e rheometer with a 50mm roughened cone plate tool, we determine the physical properties of different 3D microgel matrices. By unidirectionally shearing the samples at varying shear rates and recording the consequent shear responses as well as viscosities, we find that the material exhibits shear-dependent rheological properties as well as yield stress behaviour — i.e., above a threshold shear rate, the material behaves like a fluid (shear stress has power law dependence on shear rate), whereas below the threshold shear rate, the material behaves like a soft solid (shear stress is independent of shear rate) ^35^. We additionally apply the Herschel-Bulkley model for yield stress fluids to determine the characteristic yield stress and associated crossover shear rates for each of these microgel matrices (**Supplementary Table 1**). We also report the bulk stiffnesses of these matrices by subjecting them to small amplitude (~1%) oscillatory shear strain over varying frequencies and record the resultant elastic storage modulus (*G’*, indicating solid-like behaviour) as well as viscous loss modulus (*G’’*, indicating fluid-like behaviour). From this, we also infer that the rheological properties of our 3D matrix remain conserved irrespective of the time period over which deforming shear is applied — indicating time-independent material properties.

### 4.3. Nematode culturing protocols

*Caenorhabditis elegans* strains used in this study were sourced from the *Caenorhabditis* Genetics Centre (CGC, University of Minnesota, Minneapolis). Specimen denoted as the wild-type belongs to the N2 strain (Bristol variety), while the soft touch-insensitive mutant *mec-4* is strain CB1229 [genotype: mec-4(e1339) X]. Maintenance cultures of nematodes are carried out on NGM (nematode growth medium) agar plates (2.5% w/v agar), which are seeded with the *E. coli* strain OP50 (a uracil auxotroph) as a food source. Plates with nematodes are maintained at 21°C, with regular transfers to fresh plates such as to prevent starvation and overcrowding-induced stresses. The M9-based nutrient broth comprises 0.6% (w/v) disodium hydrogen phosphate, 0.3 (w/v%) monopotassium phosphate, 0.5% sodium chloride and 0.1% ammonium chloride dissolved in double-distilled water and autoclaved before use. Twenty-four hours before all assays, we transfer healthy nematodes at late L4-stage to fresh NGM agar plates and allow them to grow into adult nematodes with a body length of approximately 1mm ± 100 µm. Approximately 4-5 nematodes per assay (to minimize the interactions among the nematodes) are first washed using the respective yield stress gel to remove possible bacterial coatings from the nematode’s body surface and eliminate chemotactic responses, following which these are transferred to the imaging setup containing 50-100 µL of the 3D microgel matrix, wherein we allow for a 15-minute incubation to acclimatize the nematodes. To visualize the flow field around a nematode in motion, we uniformly disperse 10-micron diameter polystyrene green fluorescent beads in a microgel matrix and add nematodes to this before imaging.

### 4.4. Imaging setup

For all image acquisitions, we use a Nikon Eclipse Ti2 point-scanning laser confocal microscope. Assay setups are prepared by glueing together two transparent 100 micron thick A4 sheets with UV-curable Norland Optical Adhesive 81, to generate 200 micron thick slabs. Two circular 1 cm diameter holes are punched into this slab, to one side of which we affix a cover glass slide. This generates a confined space of <100 µL within which the nematode-laden microgel mix is loaded, and the chamber is sealed with a cover glass with excess gel being scrapped off. Images are acquired using a 4X objective ~20 min after loading inside the imaging chamber, for a period of 1-2 minutes per image, and all experiments are conducted within approximately 10-15 minutes. At least twenty nematodes are analysed for all experimental conditions, across both strain types.

### 4.5. Quantitative image analysis

Brightfield images from all timelapse data are processed using ImageJ. Using background subtraction methods, we obtain binary images which show the nematode body in motion through the microgel matrix. From these, we separate out frames capturing the undulatory motion of individual nematodes, over time windows within which they execute at least one complete sinusoidal motion. Idealizing the nematode body as a sinusoidal wave, we measure the amplitude, wavelength, and frequency of motion as outlined in **Figure 2b**. Subsequently, we obtain the characteristic time period of motion for different degrees of physical confinement. Using custom MATLAB scripts, we track the nematode body centroid, from which we measure the mean square displacement (MSD) as:

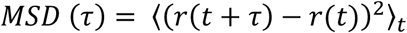

Where, *r* is position and τ is lag time. The root mean square speed of forward motion is calculated from the MSD as:

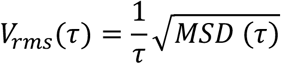

We then compare the characteristic time period of motion as measured by manual image analysis (described above) and find the root mean square speed most closely corresponding to this time interval, which we assign as the *V*_*rms*_ for each particular microgel matrix.

### 4.6. Statistical analysis

In general, for any term *h*(*x, y*) that comprises two independent variables *x* and *y* each with their own mean values and errors (*dx* and *dy*), the propagated error of the overall term (*dh*) can be written as

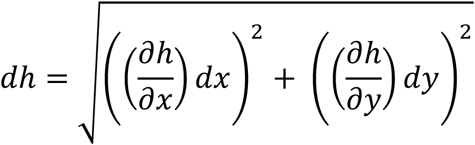

For a general quantity

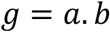

the propagated error *dg* will be as

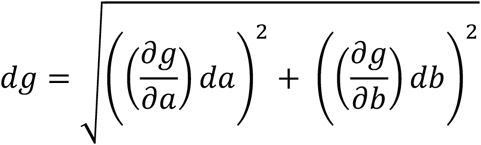

which reduces to

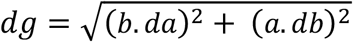

Multiplying and dividing the R.H.S. by *g*, we get

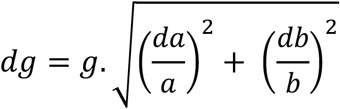

Similarly, if

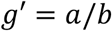

the propagated error *dg* will be as

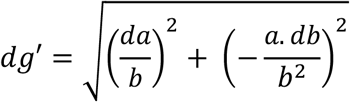

Multiplying and dividing the R.H.S. by *g*′, we get

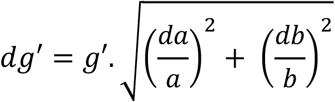

Using this, we calculate the propagated error for each of *A*/*λ, f*· A, *f*· λ, 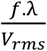, and 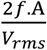 as:

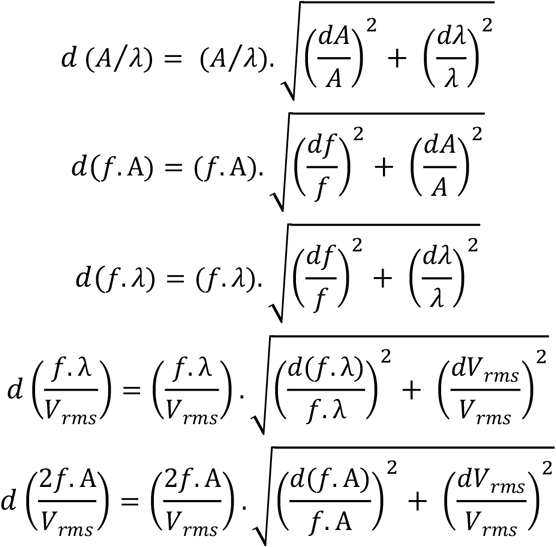

Experimentally, we define the Bingham number as

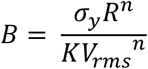

Obtaining a partial derivative for each term in this, we have

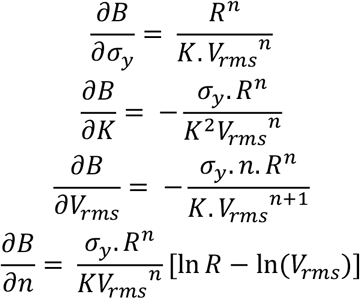

Using the formula for propagated error, we have

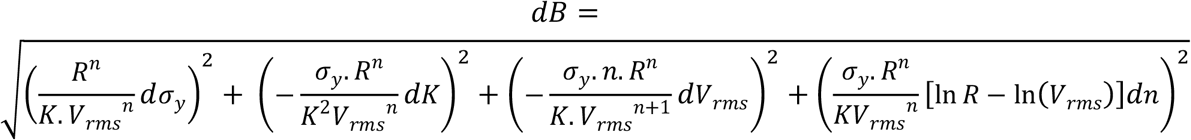

This can be simplified as

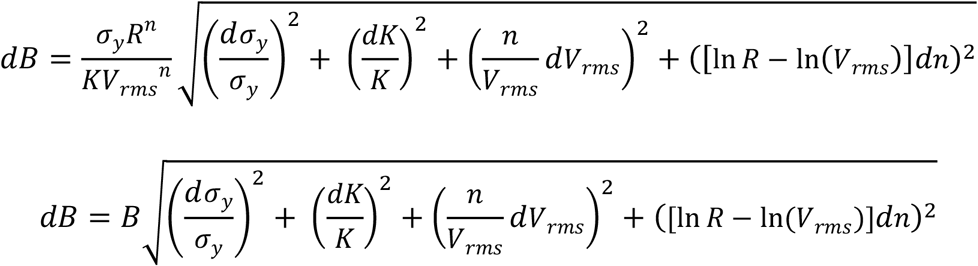

For calculating the crossover shear rate, 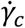, we refer to its definition as

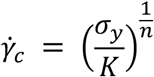

The partial derivatives for all terms on the R.H.S. will be as

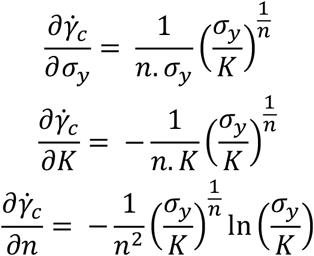

Hence, the propagated error 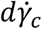will be as:

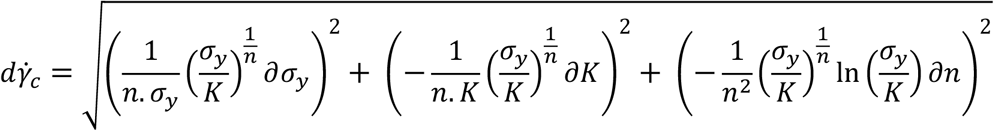

which further reduces to

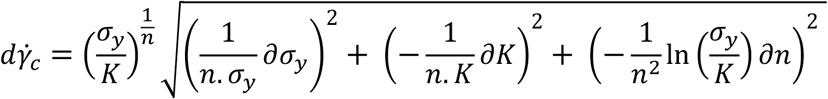

Finally, we calculate the *T*^∗^ as

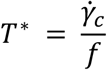

wherein, the propagated error is given as

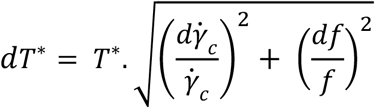

## Acknowledgements

S.D. acknowledges the Graduate Trainee program at NCBS. M.S. and A.B.A. acknowledge NCBS PhD program for funding support. S.K. acknowledges the DISHA scholarship from NISER. T.B. acknowledges NCBS intramural funding for research support. We thank Dr. Abhishek Bhattacharyya, Prof. Anindya Ghosh Roy, and Prof. Kabita Babu for sharing *C. elegans* strains. We are grateful to Dr. Abhishek Bhattacharyya and Ms. Selvanayaki E for the user support at the *C. elegans* facility and for training the authors on *C. elegans* culture protocols, as well as Ms. Sneha Hegde for helping with the assay design. We acknowledge several valuable inputs from discussions with Prof. Madan Rao. We also thank J.P. Baneerjee for his contributions towards initial efforts at simulating nematode motion in granular baths. We are also thankful to Prof. Paulo Arratia and Dr. David Gangon for insightful discussions as well as for sharing the NABAS software system.

## Author contributions

T.B. conceived, designed, and supervised the study. S.D. performed the majority of the experiments, designed the imaging setup, and optimized all protocols. S.K., A.B.A., and M.S. collected selected subsets of experimental data. D.H. conceptualized the fundamental framework underlying the theoretical model of nematode motion, with subsequent modifications from T.B. and M.S. towards the final formulation presented in this work. M.S. and S.K. with guidance from T.B. performed the formal data analysis. S.P.K. contributed towards designing key experimental protocols and participated in critical discussions. S.T. contributed towards the primary assay design, provided infrastructural support for preliminary experiments by S.D., and provided critical inputs towards interpreting the experimental observations. M.S. curated all data and prepared all figures. M.S. and T.B. wrote the manuscript, with support from D.H. Funding support for this project was acquired by T.B.

## Supplementary Information

### Section 1. Supplementary Data

**Supplementary Table 1:**
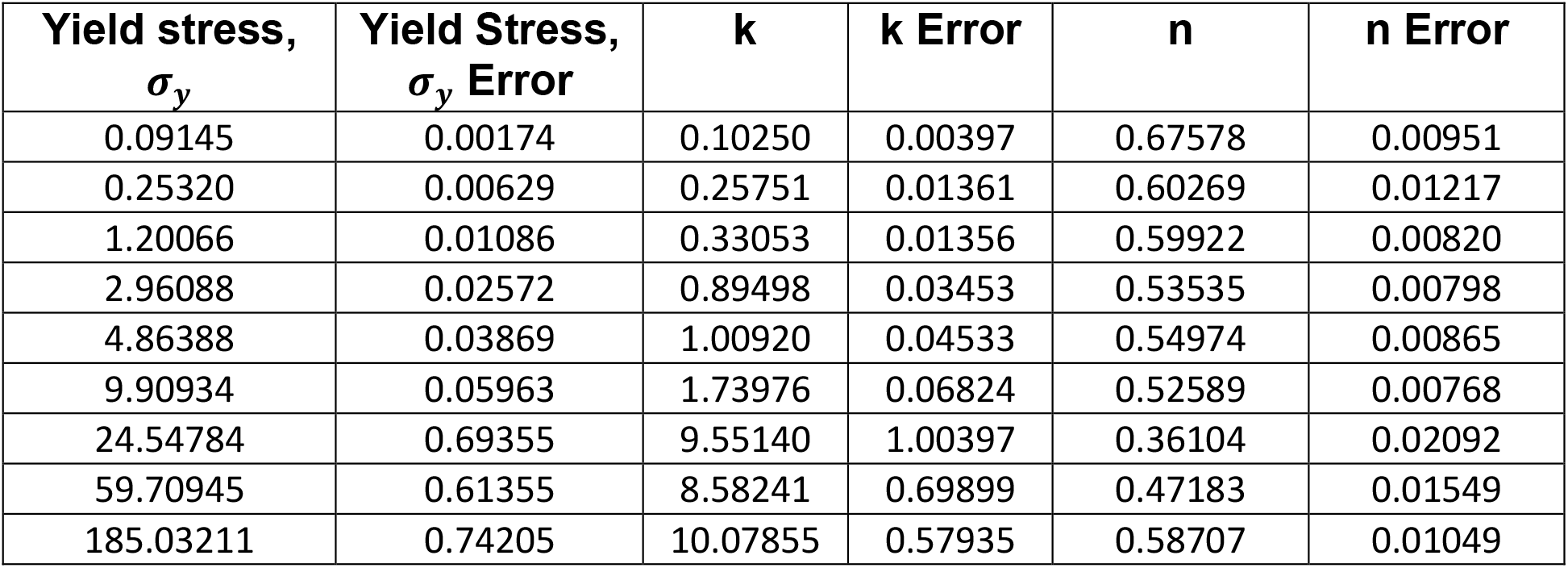
Rheological fitting parameters for the 3D microgel growth media used in this study.

| **Yield stress,**<br>**$\sigma_y$** | **Yield Stress,**<br>**$\sigma_y$ Error** | **k** | **k Error** | **n** | **n Error** |
| --- | --- | --- | --- | --- | --- |
| 0.09145 | 0.00174 | 0.10250 | 0.00397 | 0.67578 | 0.00951 |
| 0.25320 | 0.00629 | 0.25751 | 0.01361 | 0.60269 | 0.01217 |
| 1.20066 | 0.01086 | 0.33053 | 0.01356 | 0.59922 | 0.00820 |
| 2.96088 | 0.02572 | 0.89498 | 0.03453 | 0.53535 | 0.00798 |
| 4.86388 | 0.03869 | 1.00920 | 0.04533 | 0.54974 | 0.00865 |
| 9.90934 | 0.05963 | 1.73976 | 0.06824 | 0.52589 | 0.00768 |
| 24.54784 | 0.69355 | 9.55140 | 1.00397 | 0.36104 | 0.02092 |
| 59.70945 | 0.61355 | 8.58241 | 0.69899 | 0.47183 | 0.01549 |
| 185.03211 | 0.74205 | 10.07855 | 0.57935 | 0.58707 | 0.01049 |

**Supplementary Figure 1:**
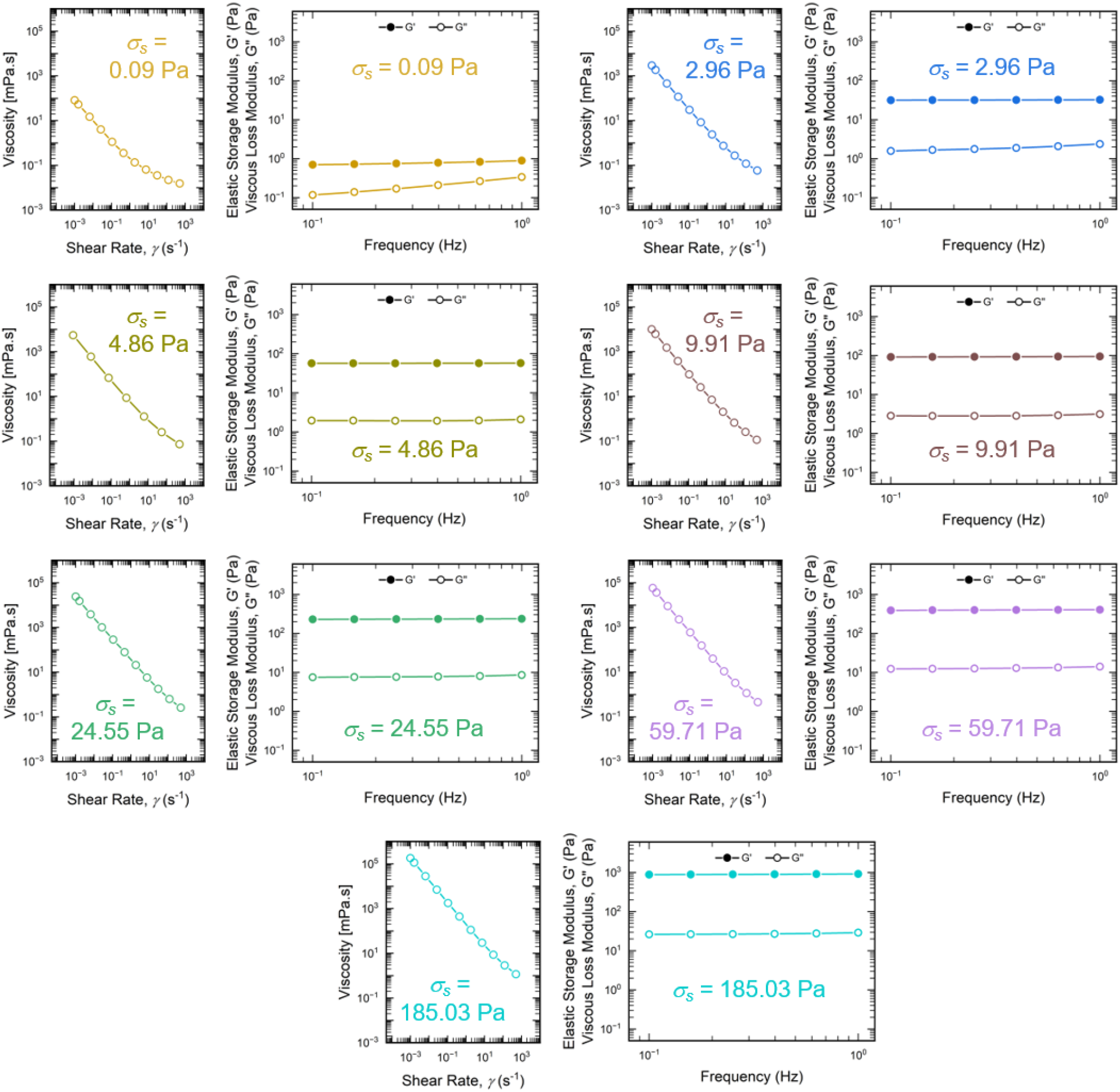
The mechanically tunable properties of the 3D matrix are characterized by shear rheology. Corresponding to the different yield stress properties of these matrices, we also find varying viscosities and varying bulk stiffnesses, showing how the system can be engineered to realize different degrees of physical confinement.

**Supplementary Figure 2:**
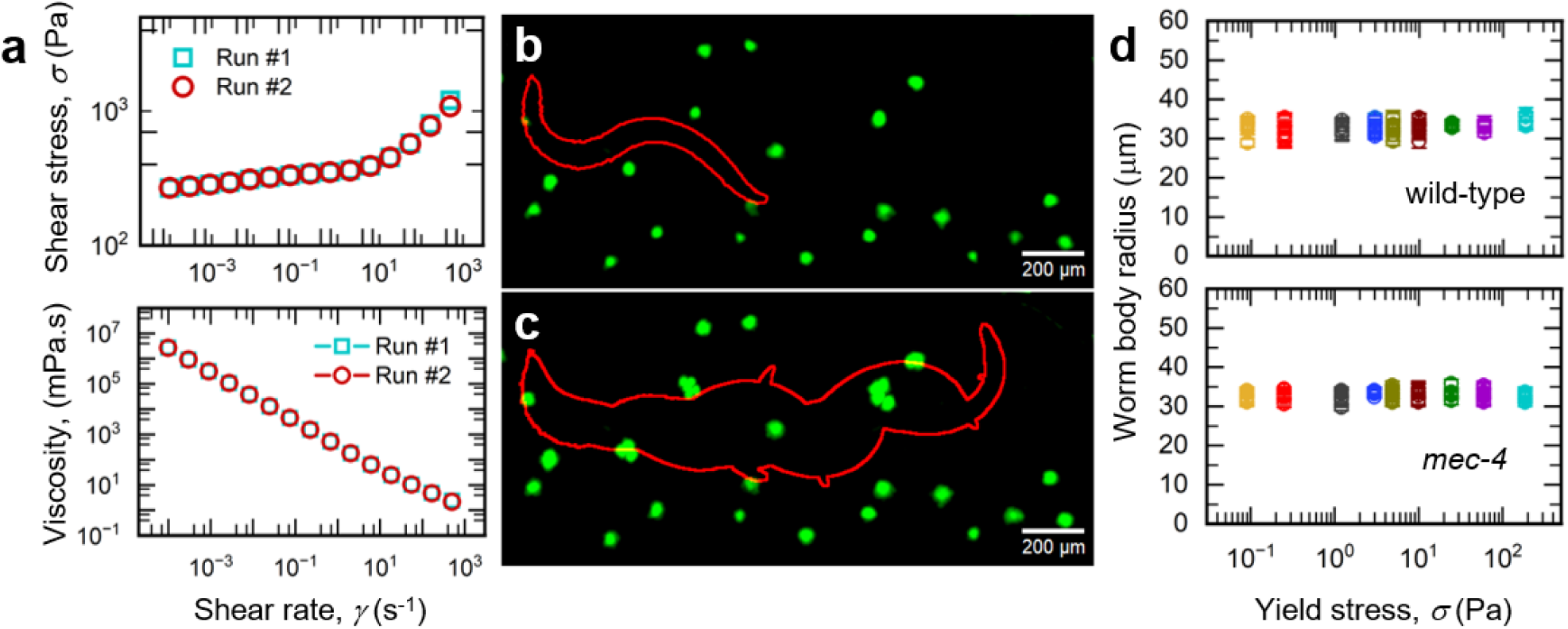
(a) Consecutive rounds of unidirectional shear over a range of shear rates do not alter the rheological properties of the 3D microgel media, illustrating its self-healing nature following mechanical shear. (b) By visualizing the nematode motion in a 3D matrix laden with green fluorescent beads, we find that the nematode (outlined in red) only deforms the matrix in a localized manner in the region immediately surrounding its body. (c) At locations distant from the nematode body — indicated by beads that fall well outside the region swept by the nematode motion (outlined in red) — no direct effect of the nematode’s movement are discernible, indicating that the motion-induced shear effects do not contribute towards a global deformation of the jammed microgel matrix. (d) Confinement within 3D microgel growth media does not induce aberrant morphologies by mechanically deforming or compressing the nematode bodies.

**Supplementary Figure 3:**
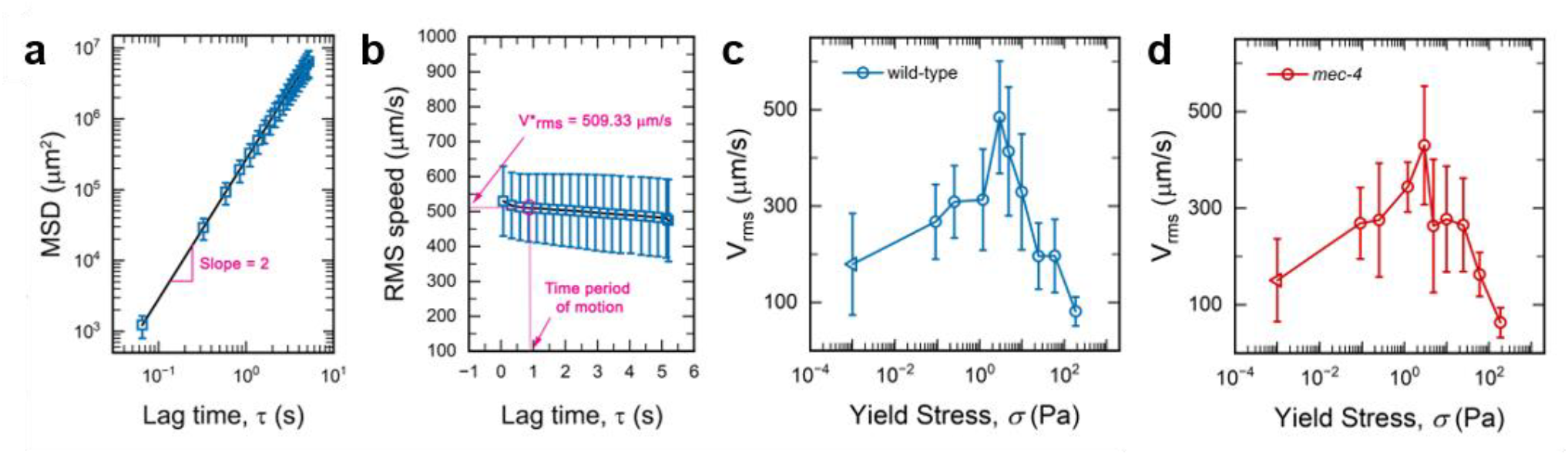
(a) Tracking nematode motion within the 3D yield stress matrix reveals that nematodes follow a ballistic motion over short time scales. (b) By determining the characteristic time periods of motion from analysis of the undulatory motion, we find the nematode’s forward propulsive speed, as given by the rate of displacement of the body centroid. (c and d) The non-monotonic trend in forward propulsive speed remains conserved for both wild-type and mechanosensory mutant strains regardless of the time period of motion chosen for analysis, as evidenced by taking an average speed of the nematodes across all time periods over the entire duration of tracking. In all the plots of panel c and d, the triangular symbols represent data from nematodes moving through free liquid. Since free liquid does not have a yield stress, an arbitrary value of 10^−3^ Pa (viscosity of water times one second) is assigned to include these data on the same plot.

**Supplementary Figure 4:**
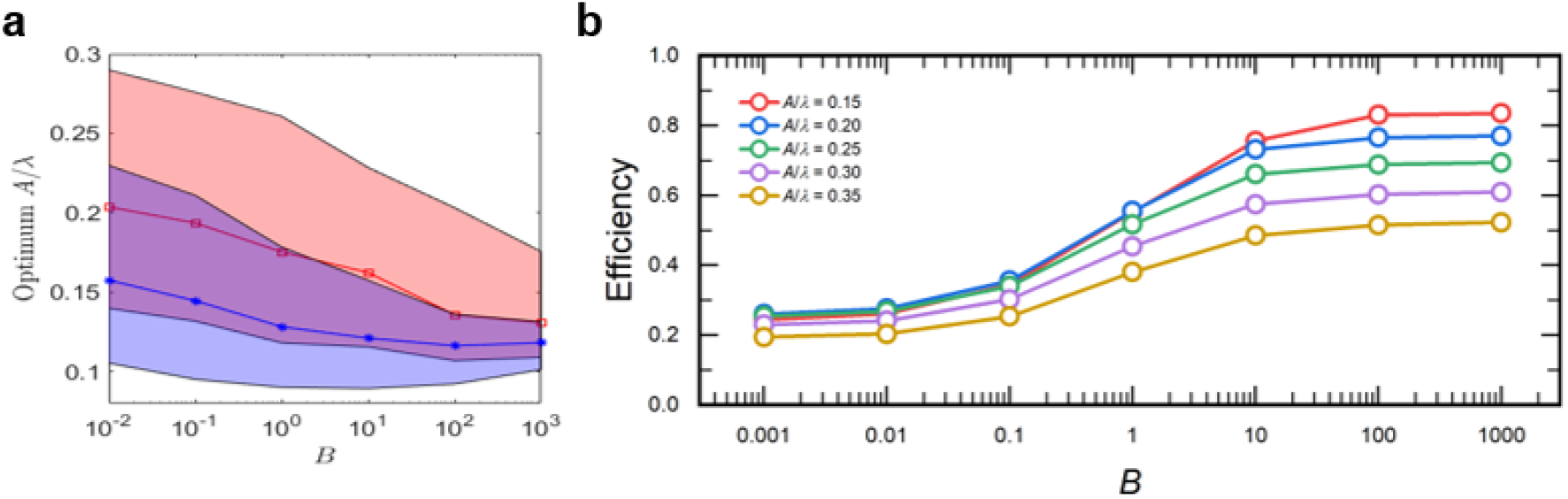
(a) Using numerical analysis, we screen for the optimum *A*/*λ* ratio across different mechanical regimes (*B*) and find this to be of the order of 0.1 ≤ *A*/*λ* < 0.5. Analysing the Lighthill efficiency across different mechanical regimes reveals a generalized plateauing behaviour when *B* > 1 (solid-like behaviour) for all body forms (*A*/*λ*).

**Supplementary Figure 5:**
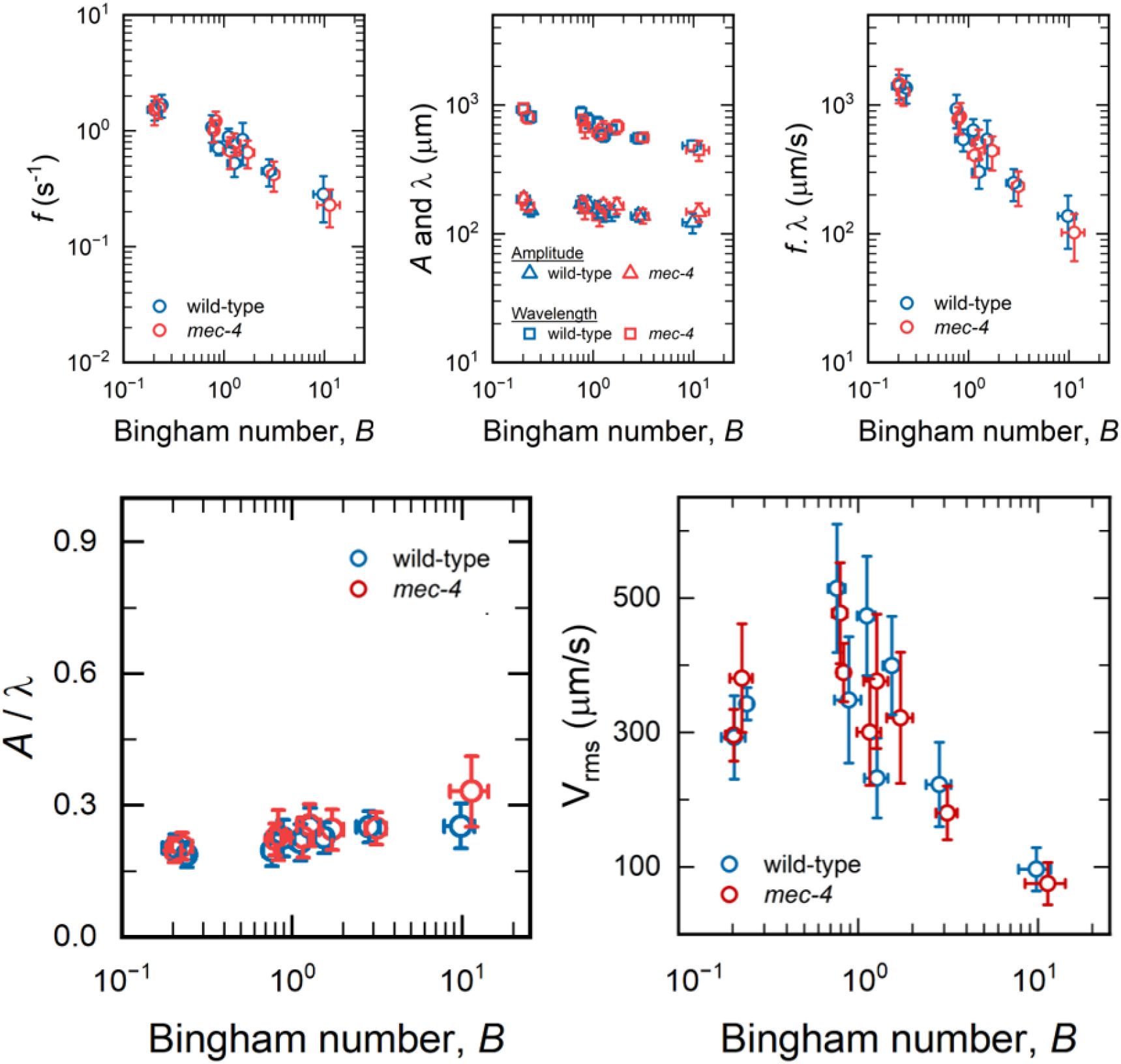
Parameters describing the nematode motion, plotted as a function of the experimentally calculated Bingham number.

**Supplementary Figure 6:**
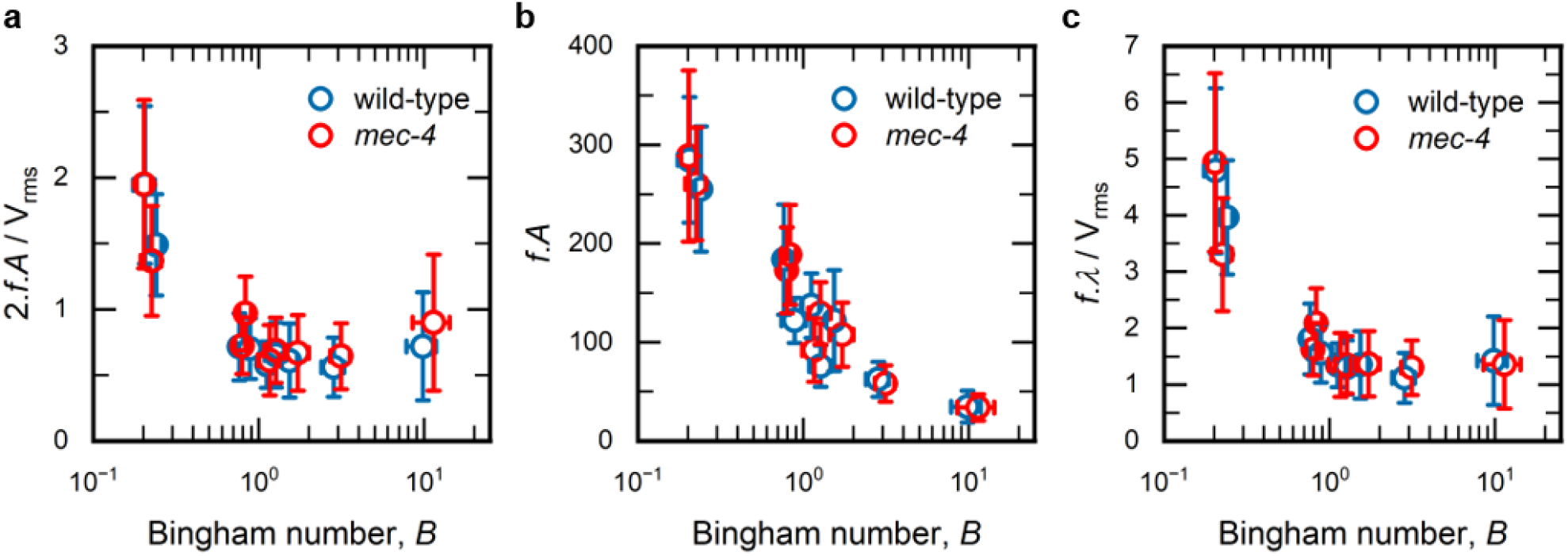
Analysis of the (a) Strouhal number (b) sideways component of motion and (c) ratio between wave speed and propulsive speed across different values of the Bingham number calculated experimentally.

**Supplementary Figure 7:**
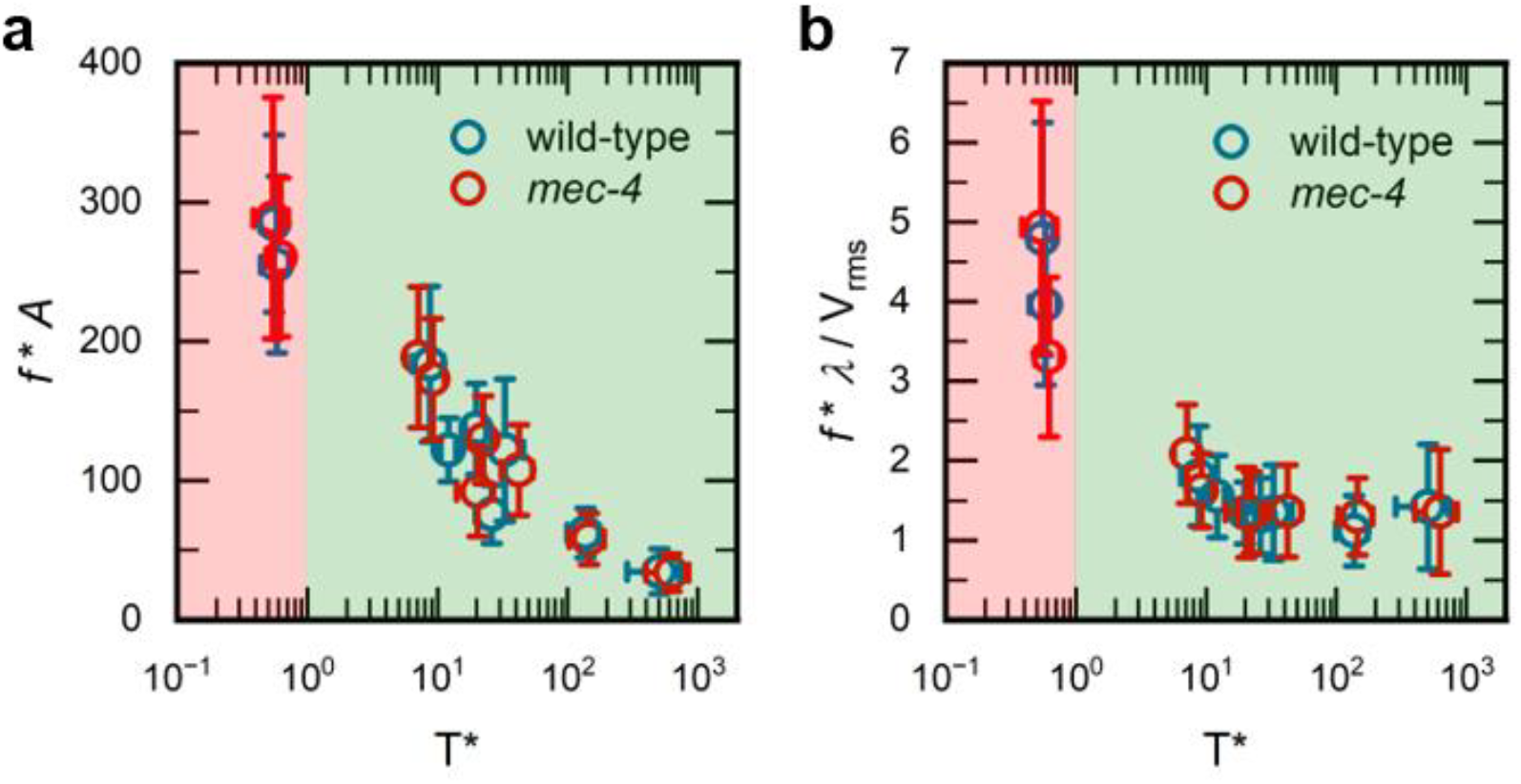
We show that both (a) the sideways component of motion and (b) ratio between wave speed and propulsive speed show distinct behaviours going from fluid-like (*T\** < 1) to solid-like (*T\** > 1) regimes. While the sideways motion decreases with increasing physical confinement, we find that the wave speed along nematode bodies matches the propulsive speed under increasing physical confinement.

### Section 2. Theoretical formulation of a microswimmer in a yield stress fluid

Given the yield stress nature of the 3D microgel matrix used in our study, we can construct a material framework for this based on the Herschel-Bulkley model for yield stress fluids. The key considerations are that such materials show reversible, shear-dependent, fluid-to-solid transition based on the shear stress imparted by the nematode’s motion through the matrix. Hence, we can model the deviatoric stress within such a Herschel-Bulkley-like fluid as

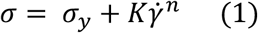

for shear stress σ, yield stress σ_*y*_, consistency *K*, index *n*, and the strain rate 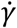. We can then consider a nematode in motion through this matrix as an inextensible cylindrical swimmer of radius R, that propagates in a sinusoidal fashion following the waveform

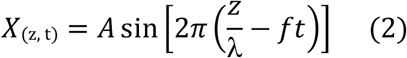

wherein amplitude *A*, wavelength *λ*, wave speed *c*, and frequency *f* = *c*/*λ*, represent the sinusoidal propulsion pattern.

Next, to model the effect of forces associated with motion through the yield stress fluid on the nematode body, we invoke assumptions from resistive force theory. Principally, we consider that the global effect of all forces on the nematode can be captured by considering the effects of these on each infinitesimal element of the worm body. This also brings in the asymptotic condition that the length scale across which we will study the motion is much longer than the radius of the nematode. This allows us to break down the overall nematode body into a collection of isolated filaments, each being acted upon by the set of forces and thereby attaining a resultant orientation and speed, while remaining coupled to the ensemble structure. Importantly, this assumption is particularly well-suited for a Herschel-Bulkley fluid, as opposed to a Newtonian fluid, since the yield stress nature of the matrix limits the extent to which flow-induced shear can deform the nematode body. While prior treatments of undulatory swimmers by such variations of slender body theory assume an agent capable of generating infinite power, herein, we stay close to the experimental paradigm by considering that over the course of motion, a nematode generates a finite amount of power to navigate through the 3D matrix. Accordingly, the power expended per unit length can be expressed as

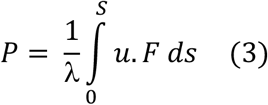

where for each finite element *ds* of the body length, *u* is the local velocity, *F* is the force on the swimmer’s body, and the overall integral is over the path of the nematode’s body for one wavelength.

However, we note that our biophysical measurements of the nematode motion cannot explicitly capture its power generation. Rather, our observations are based on the propulsive behaviour itself, as captured by the components describing its sinusoidal motion. Hence, it becomes necessary to explore a relation that directly links power generation to quantifiable parameters of the nematode’s motion, which are also dependent on the degree of physical confinement.

Accordingly, we note that the power term *P* has dimensions similar to the product of force and velocity. The force component here can now be related to the deviatoric stress relation imposed by the Herschel-Bulkley fluid-like nature of the 3D matrix under nematode motion-induced shear, by introducing some velocity scale V, such that we obtain

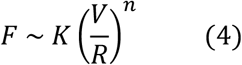

which, considering that power is the product of force and velocity, becomes

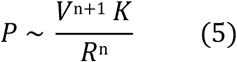

which itself interchangeably becomes (by fixing the power generated by the swimmer as a constant over the period of motion)

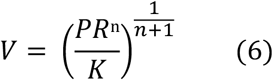

These relations now allow us to describe the power generation in terms more amenable to our experimental system — while both *V* and *R* can be measured by measuring the nematode motion, the parameter *K* can be derived by fitting the Herschel-Bulkley equation to the shear stress response of the 3D microgel matrix across varying shear rates.

Further, considering that we are dealing with motion through a yield stress material, we can define a dimensionless formulation of the yield stress itself, termed the Bingham number, which takes into account both the material properties of the matrix (in terms of yield stress σ_*y*_, consistency *K*, and index *n*) as well as appropriate descriptors of the nematode motion (in terms of radius *R* and velocity *V*), as follows:

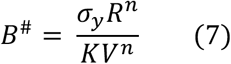

We can also express this in terms of the power generated by the nematode, using the velocity scale term from (6) above, as

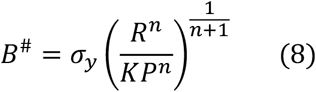

We note that while *B*^#^ and P give an ensemble idea of the yield stress nature of the bath and power generated by the nematode’s motion through the 3D matrix, it is also necessary to better describe the local interactions that govern how the nematode attempts to yield this Herschel-Bulkley fluid-like matrix to achieve motion. Hence, we define both a rescaled Bingham number 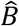, and a rescaled power generation term 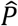, in order to capture these events. Using previous results^27^, we can obtain exact expressions for both 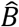 and 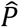, which also account for the wave speed (c) of the nematode’s undulatory motion. These are as follow

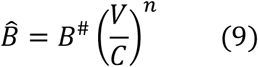

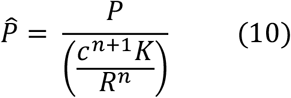

Note that *B*^#^corresponds to the Bingham number representation, *B*, as used in the main text for plotting the numerical simulations data.

It is also useful to obtain a handier expression for the rescaled local 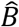 and 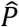, in terms of quantities that may be readily inferred from direct biophysical measurements. Hence, we obtain a direct relation between 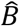 and 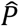 in terms of the dimensionless Bingham number *B*^#^ by effecting appropriate substitutions from the aforementioned relations, which is as

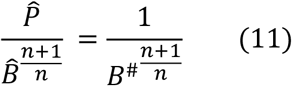

Rewriting this, we find

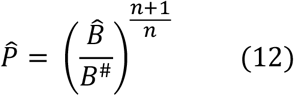

This expression enables a direct comparison with previous results^27^, which also reported a numerical expression for 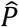 with the form

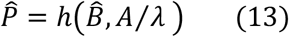

Using these, we can now redefine the function *h* in terms of *B*^#^ and 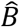 as

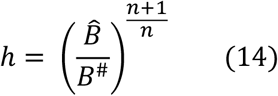

While there does not exist an analytical expression for *h*, from prior work, we can find approximations at the asymptotic limits which hold for the condition wherein large 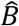 dominates (representing high yield stress regimes — precisely where we experimentally observe a decay in the nematode’s propulsive speed). Such an asymptotic form of *h* is given by

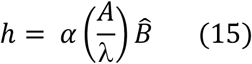

for some constant *α* and the condition wherein *A*/*λ* ≤ 1, i.e., small values of *A*/*λ*. This condition is contingent on the worm prioritizing forward motion while minimizing the sideways motion (small amplitude movement), which is an expected outcome in high yield stress regimes, wherein increased power generation is likely required to accomplish motion. Hence, it is a likely assumption that considering a worm that can generate only a finite amount of power, it will likely attempt to minimize wasteful sideways displacements, and instead, optimize for forward motion — thereby, entering a propulsive regime characterized by a small value of *A*/*λ*.

To obtain a tractable solution for these expressions, we now equate (14) and (15), which gives

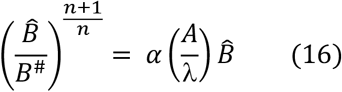

Following rearrangements between the terms, we get

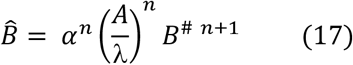

Separately, using the previous definition of 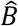 from (9), we can obtain the wave speed *c* of the nematode’s sinusoidal motion in the following form

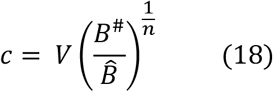

We can now replace 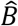 in (18), with the formulation in (17), to obtain

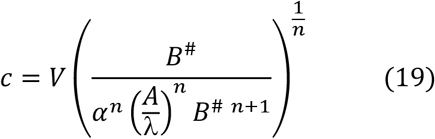

which then simplifies into

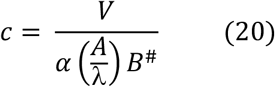

finally allowing us to resolve *B*^#^ into its constituent terms from (7), to get

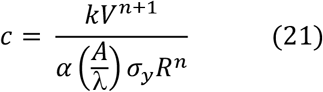

However, this velocity scale *V* still remains an incompletely characterized quantity. Hence, we use the relation from (5), to rewrite this in terms of the power *P* as

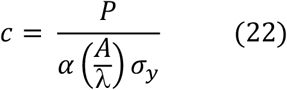

Considering that we have already defined power as the product of force and velocity, we can elementarily work out the following relations

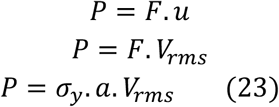

Herein, we can now define the local velocity *u* using the experimentally determined V_rms_, as well as rewrite the force *F* in terms of the yield stress *τ*_*y*_ and cross-sectional area *a*. Critically, our experimental data shows that

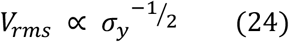

Relating these using some factor *δ*, we can now rewrite (23) as

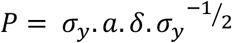

which then resolves to

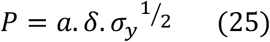

Finally, replacing this formulation of *P* in (22), we obtain

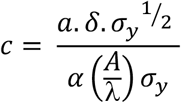

which yields the relation

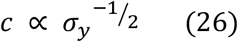

This prediction implies that the effects of increasingly solid-like regimes (characterized by *B*^#^ >> 1) on nematode motion manifest as a power law dependence of the worm’s wave speed on the matrix yield stress, following the scaling stated in (26). It should be noted that this relation is also contingent on the body form, represented by the ratio of amplitude and wavelength of the nematode body’s undulations, being actively maintained at a constant and small magnitude (*A*/*λ* ≤ 1) under high yield stress regimes, ensuring minimal wastage of power and optimization for forward motion. It is conceivable that this alteration to the undulatory motion is key towards ensuring efficient motion with an optimal expenditure of power generation under higher degrees of physical confinement, such as may be evaluated by comparing the magnitude of sideways motion (product of frequency and amplitude) with the magnitude of forward propulsive motion (*V*_*rms*_) — the Strouhal number.

## References

1. Félix, M. A. & Braendle, C. The natural history of Caenorhabditis elegans. Current Biology 20, R965–R969 (2010).

2. Schulenburg, H. & Félix, M. A. The Natural Biotic Environment of Caenorhabditis elegans. Genetics 206, 55–86 (2017).

3. Xu, Y. et al. Dopamine receptor DOP-1 engages a sleep pathway to modulate swimming in C. elegans. iScience 24, 102247 (2021).

4. Vidal-Gadea, A. et al. Caenorhabditis elegans selects distinct crawling and swimming gaits via dopamine and serotonin. Proc Natl Acad Sci U S A 108, 17504–9 (2011).

5. Ghosh, R. & Emmons, S. W. Episodic swimming behavior in the nematode C. elegans. Journal of Experimental Biology 211, 3703–3711 (2008).

6. Laranjeiro, R., Harinath, G., Burke, D., Braeckman, B. P. & Driscoll, M. Single swim sessions in C. elegans induce key features of mammalian exercise. BMC Biol 15, 30 (2017).

7. Pierce-Shimomura, J. T. et al. Genetic analysis of crawling and swimming locomotory patterns in C. elegans. Proc Natl Acad Sci U S A 105, 20982 (2008).

8. Lebois, F. et al. Locomotion Control of Caenorhabditis elegans through Confinement. Biophys J 102, 2791 (2012).

9. Korta, J., Clark, D. A., Gabel, C. V., Mahadevan, L. & Samuel, A. D. T. Mechanosensation and mechanical load modulate the locomotory gait of swimming C. elegans. J Exp Biol 210, 2383–2389 (2007).

10. Beron, C. et al. The burrowing behavior of the nematode Caenorhabditis elegans: a new assay for the study of neuromuscular disorders. Genes Brain Behav 14, 357–368 (2015).

11. Shen, X. N. & Arratia, P. E. Undulatory swimming in viscoelastic fluids. Phys Rev Lett 106, 208101 (2011).

12. Gagnon, D. A., Shen, X. N. & Arratia, P. E. Undulatory swimming in fluids with polymer networks. Europhys Lett 104, 14004 (2013).

13. Sznitman, J. & Arratia, P. E. Locomotion Through Complex Fluids: An Experimental View. 245–281 (2015) doi:10.1007/978-1-4939-2065-5_7.

14. Park, S. et al. Enhanced Caenorhabditis elegans Locomotion in a Structured Microfluidic Environment. PLoS One 3, e2550 (2008).

15. Gagnon, D. A. & Arratia, P. E. The cost of swimming in generalized Newtonian fluids: experiments with C. elegans. J Fluid Mech 800, 753–765 (2016).

16. Sznitman, J., Shen, X., Purohit, P. K. & Arratia, P. E. The effects of fluid viscosity on the kinematics and material properties of C. elegans swimming at low reynolds number. Proceedings of the Society for Experimental Mechanics, Inc. 67, 1303–1311 (2010).

17. Sznitman, J., Purohit, P. K., Krajacic, P., Lamitina, T. & Arratia, P. E. Material Properties of Caenorhabditis elegans Swimming at Low Reynolds Number. Biophys J 98, 617 (2010).

18. Park, J. S., Kim, D., Shin, J. H. & Weitz, D. A. Efficient nematode swimming in a shear thinning colloidal suspension. Soft Matter 12, 1892–1897 (2016).

19. Guisnet, A., Maitra, M., Pradhan, S. & Hendricks, M. A three-dimensional habitat for C. elegans environmental enrichment. PLoS One 16, e0245139 (2021).

20. Tahernia, M., Mohammadifar, M. & Choi, S. Paper-Supported High-Throughput 3D Culturing, Trapping, and Monitoring of Caenorhabditis Elegans. Micromachines (Basel) 11, (2020).

21. Jung, S. Caenorhabditis elegans swimming in a saturated particulate system. Physics of Fluids 22, 3–6 (2010).

22. Juarez, G., Lu, K., Sznitman, J. & Arratia, P. E. Motility of small nematodes in wet granular media. Europhys Lett 92, 44002 (2010).

23. Sreepadmanabh, M., Arun, A. B. & Bhattacharjee, T. Design approaches for 3D cell culture and 3D bioprinting platforms. Biophys Rev 5, (2024).

24. Bhattacharjee, T. et al. Liquid-like Solids Support Cells in 3D. ACS Biomater Sci Eng 2, 1787–1795 (2016).

25. Bhattacharjee, T. et al. Polyelectrolyte scaling laws for microgel yielding near jamming. Soft Matter 14, 1559–1570 (2018).

26. O’Hagan, R., Chalfie, M. & Goodman, M. B. The MEC-4 DEG/ENaC channel of Caenorhabditis elegans touch receptor neurons transduces mechanical signals. Nature Neuroscience 2004 8:1 8, 43–50 (2004).

27. Hewitt, D. R. & Balmforth, N. J. Locomotion with a wavy cylindrical filament in a yield-stress fluid. J Fluid Mech 936, A17 (2022).

28. Hewitt, D. R. Swimming in viscoplastic fluids. Rheol Acta 63, 673–688 (2024).

29. Kudrolli, A. & Ramirez, B. Burrowing dynamics of aquatic worms in soft sediments. Proc Natl Acad Sci U S A 116, 25569–25574 (2019).

30. Dorgan, K. M., Law, C. J. & Rouse, G. W. Meandering worms: mechanics of undulatory burrowing in muds. Proceedings of the Royal Society B: Biological Sciences 280, (2013).

31. Ahamed, T., Costa, A. C. & Stephens, G. J. Capturing the continuous complexity of behaviour in Caenorhabditis elegans. Nature Physics 2020 17:2 17, 275–283 (2020).

32. Tytell, E. D., Holmes, P. & Cohen, A. H. Spikes alone do not behavior make: why neuroscience needs biomechanics. Curr Opin Neurobiol 21, 816–822 (2011).

33. Sreepadmanabh, M. et al. Cell shape affects bacterial colony growth under physical confinement. Nature Communications 2024 15:1 15, 1–13 (2024).

34. Bhattacharjee, T. & Datta, S. S. Bacterial hopping and trapping in porous media. Nature Communications 2019 10:1 10, 1–9 (2019).

35. Sreepadmanabh, M., Ganesh, M., Bhat, R. & Bhattacharjee, T. Jammed microgel growth medium prepared by flash-solidification of agarose for 3D cell culture and 3D bioprinting. Biomedical Materials 18, 045011 (2023).

